# Repeat mediated excision of gene drive elements for restoring wild-type populations

**DOI:** 10.1101/2023.11.23.568397

**Authors:** Pratima R Chennuri, Josef Zapletal, Raquel D Monfardini, Martial Loth Ndeffo-Mbah, Zach N Adelman, Kevin M Myles

## Abstract

We demonstrate here that single strand annealing (SSA) repair can be co-opted for the precise autocatalytic excision of a drive element. Although SSA is not the predominant form of DNA repair in eukaryotic organisms, we increased the likelihood of its use by engineering direct repeats at sites flanking the drive allele, and then introducing a double-strand DNA break (DSB) at a second endonuclease target site encoded within the drive allele. We have termed this technology <u>Re</u>peat <u>M</u>ediated <u>E</u>xcision of a <u>D</u>rive <u>E</u>lement (ReMEDE). Incorporation of ReMEDE into the previously described mutagenic chain reaction (MCR) gene drive, targeting the *yellow* gene of *Drosophila melanogaster*, replaced drive alleles with wild-type alleles demonstrating proof-of-principle. Although the ReMEDE system requires further research and development, the technology has a number of attractive features as a gene drive mitigation strategy, chief among these the potential to restore a wild-type population without releasing additional transgenic organisms or large-scale environmental engineering efforts.

## Introduction

Gene drives are currently being developed as tools for the control of sexually reproducing populations of organisms that negatively impact public health, agriculture, and conservation efforts (Burt, 2003; Esvelt et al., 2014). The core technology underlying various gene drive-based control methods currently under development are transgenes that will be inherited at super-Mendelian rates, after the individuals that have been engineered to carry them mate with wild populations. Examples of organisms that are currently being considered for control with this technology include insects that adversely affect public health, e.g., mosquito vector species (Gantz and Bier, 2015; Hammond et al., 2016; Kyrou et al., 2018), agriculture, e.g., pest species (Courtier-Orgogozo et al., 2017; Scott et al., 2018); and invasive rodents threatening the natural fauna and flora of island ecosystems, e.g., rat and mouse species (Grunwald et al., 2019; Leitschuh et al., 2018). Although the concept of large-scale genetic engineering projects to control or modify entire populations has existed for some time (Curtis, 1968, p. 19; Deredec et al., 2008), the description and development of the type II CRISPR/Cas9 system for introducing targeted double-strand DNA breaks (DSBs) (Jinek et al., 2012), with subsequent demonstration of the system’s potential for inducing mutagenic chain reactions (Gantz and Bier, 2015), has spurred a period of rapid technological development in this area. In theory, the CRISPR/Cas9 system is capable of cutting the DNA of any organism. Thus, many different gene drive configurations have been developed or theorized that are based on this programmable nuclease system originally from bacteria and archaea (Esvelt et al., 2014; Raban et al., 2020).

While none of these configurations have yet been tested in the field, the development of CRISPR/Cas9-based homing gene drives are proceeding rapidly (Raban et al., 2020). In this configuration, the transgene encodes the Cas9 endonuclease and a guide RNA (gRNA). The synthetic gRNA will direct the Cas9 to a predetermined target site present in the organism’s genome, resulting in a DSB at the target location. As eukaryotic organisms routinely repair DSBs, engineering the transgene with the sequences flanking the DSB (i.e., homology arms) enables a form of homology directed repair, termed homologous recombination (HR), that is capable of restoring the integrity of the double helix structure at the break site using a nearby homologous donor sequence, usually a sister chromatid during the S and G2 phases of the cell cycle. While HR is the most common of the various homology directed repair processes, it also requires the longest lengths of sequence homology between the donor and acceptor DNA. Homology directed repair of the DSB site, with the transgene serving as the donor, results in “gene conversion”. If gene conversion occurs in the germline, the transgene will be heritable, and in theory the process repeated indefinitely in subsequent generations, converting heterozygous progeny to homozygous progeny in the germline. This process ensures that the transgenic drive allele is inherited at rates far exceeding normal ratios of Mendelian inheritance. For this reason, homing gene drives have been referred to as being “threshold-independent”, i.e., capable of being transmitted through a population following the introduction of an undefined but low number of individuals carrying the transgenic drive allele (Marshall and Akbari, 2018). However, multiple studies that have mathematically modeled the spread of homing drives predict that in reality this type of drive may only be able to spread through a population when introduced above a particular frequency, which is likely determined by a number of different factors (Alphey and Bonsall, 2014; Deredec et al., 2008; Tanaka et al., 2017; Unckless et al., 2015).

Nevertheless, the specter of introducing, at previously unimaginable scales, potentially irreversible alterations to complex ecosystems raises a number of complicated ethical, legal, and social concerns (Montenegro de Wit, 2019; NASEM, 2016). Therefore, if the potential medical, agricultural and environmental benefits of gene drive technologies are to be fully realized, these concerns will need to be mitigated. This can be achieved through the development of technologies capable of limiting, both spatially and temporally, the spread of gene drives through a population (Marshall and Akbari, 2018; Raban et al., 2020). While a number of mitigation strategies for minimizing the possibility of unanticipated harmful effects on humans, animals and the environment have been proposed (Basgall et al., 2018; DiCarlo et al., 2015; Esvelt et al., 2014; Gantz and Bier, 2016; Marshall and Akbari, 2018; Wu et al., 2016; Xu et al., 2020), current state-of-the-art technologies for limiting CRISPR/Cas9-based homing gene drives are primarily based on various designs of transgenic genetic elements (Gantz and Bier, 2016) or split drive configurations (Kandul et al., 2021; Li et al., 2020; López Del Amo et al., 2020; Oberhofer et al., 2021). A number of other systems have also been proposed, and in some cases demonstrated in laboratory studies, to limit the spread of CRISPR/Cas9-based homing gene drives (Basgall et al., 2018; Chae et al., 2020; DiCarlo et al., 2015; Esvelt et al., 2014; Taxiarchi et al., 2021). Although demonstrating the efficacy of these systems is a significant advance, it is not yet clear that any of the current technologies will be able to adequately address concerns surrounding the use of gene drives in nature. While other types of drive mechanisms also exist (Bier, 2022), and may ultimately prove to raise fewer concerns than homing drives, no drive technology has yet been deemed entirely safe and tested in nature.

Previously, we proposed a novel approach to leverage a less common form of homology directed repair, single-strand annealing (SSA), for the precise removal of gene drive alleles (Zapletal et al., 2021). In theory, this technology has the potential to permit an autonomous low-threshold drive to spread through a population over a specified period of time, after which the drive allele would be remediated from the environment, fully restoring a wild-type population. Here, we demonstrate proof-of-concept for this approach through the precise autocatalytic excision of an autonomous CRISPR/Cas9-based homing drive. This was accomplished by co-opting the SSA repair mechanism, following induction of a DSB by a second endonuclease at a target site encoded within the drive allele. Although SSA is not the predominant form of DNA repair in eukaryotic organisms, we increased the likelihood of its use by engineering direct repeats at sites flanking the drive allele. We have termed this technology <u>Re</u>peat <u>M</u>ediated <u>E</u>xcision of a <u>D</u>rive <u>E</u>lement (ReMEDE).

## Methods

### Plasmid Construction

Standard recombinant DNA techniques were used to clone all constructs. Supplementary figures S1A (yDR and yReMET), S2A (yMCR) and S2B (yReMEDE) outline plasmid construction. Complete maps and sequences are shown in Fig. S6.

### Microinjections

Plasmids were purified using the Macherey-Nagel NucleoBond Xtra Midi endotoxin-free kit (#740420) and sent to BestGene Inc. or GenetiVision for embryo microinjections. The plasmids p-yMCR.EGFP and p-yDR.EGFP were co-injected with transient sources of a single guide RNA targeting exon 2 of the *yellow* gene (pCFD3-dU6:3-y1-sgRNA) and Cas9 (pBS-Hsp70-Cas9, a gift from Melissa Harrison & Kate O’Connor-Giles & Jill Wildonger - Addgene plasmid # 46294) into a *w^1118^* stock (BDSC #3605). The plasmids p-ReMET.RFP and p-ReMEDE.RFP were co-injected with a transient source of ϕC31 into stocks carrying the construct p-yDR.EGFP. All transgenic stocks were confirmed through PCR and Sanger sequencing.

### Genomic DNA Isolation and Sequence Analysis

Genomic DNA was prepared from individual flies according to protocols from Gloor et al., 1993 (Gloor et al., 1993). PCR reactions were assembled with GoTaq®G2 Green Master Mix (#7823), purified with Zymo Research DNA Clean & Concentrator-5 kit (#4014), sequenced and analyzed with SnapGene and PolyPeakParser.

### Genetic Crosses and Phenotyping

Injected G0 transformants were crossed with *w^1118^* flies and G1 progeny screened for either EGFP or RFP positive individuals. These individuals were then outcrossed with *w^1118^* flies prior to establishing homozygous lines. The yMCR or yReMEDE males were crossed with wild-type (*w^1118^*) females in order to generate F1 females. The F1 females were then mated with wild-type males in single pair crosses for scoring F2 gene drive phenotypes.

### Population Cage Studies

Discrete small scale population studies were set up in standard fly rearing bottles (Genesee Scientific Cat. #32-130) with a design similar to those described previously (Bishop et al., 2022; Xu et al., 2020). Briefly, for each genotype an equal number of virgin females and males were seeded. All flies were removed following egg laying. The progeny from each subsequent generation were collected and then separated into two equal pools (n=150) of randomly selected flies. Flies from one pool were screened and phenotyped, while the other pool was seeded into a new cage.

### Drosophila Husbandry

The I-Site reporter line, *w^1118^; P{w^+mW.hs^=w8,I-site}7* (BDSC# 6972), was sourced from the Bloomington Drosophila Stock Center. A nos-Gal4 line, *y* w*; P{w^+mC^=GAL4-nos.NGT}40* (Kyoto DGRC# 107-748), was obtained from the Kyoto Drosophila Stock Center. Stocks were maintained on standard oatmeal/cornmeal medium at 18 °C, but all experiments were performed at 25 °C, unless otherwise noted. All gene drive lines were handled in an ACL-2 facility in accordance with standard containment protocols. All work with genetically modified organisms was performed under protocols approved by the Institutional Biosafety Committee at Texas A&M University.

## Results

### <u>Re</u>peat <u>M</u>ediated <u>E</u>xcision of a <u>T</u>ransgene (ReMET)

As an initial test of the hypothesis that the SSA repair pathway can be co-opted for the removal of gene drive alleles, we generated multiple transgenic lines containing ReMET reporter constructs flanked with SSA substrates (i.e., direct repeats) of various lengths. Briefly, the reporter constructs consist of an endonuclease, I-*Sce*I, under the control of a 5x Upstream Activation Sequence (*UASp.I-SceI*), the 18 bp I-*Sce*I target site (I-Site), and two fluorescent marker sequences, enhanced green fluorescent protein (EGFP) and red fluorescent protein (RFP), under the control of independent 3xP3 promoters. The yeast transcription activator protein, Gal4, was placed under the control of the germline-specific promoter *nanos* (*nos*) in a mifepristone (RU486)-inducible GeneSwitch expression system (Fig. 1A and S1A). Constructs were inserted into the *yellow (y)* gene of *D. melanogaster*. Expression of I-*Sce*I generates a DSB at the I-Site sequence, which facilitates SSA-based repair mediated by the presence of the direct repeats, leading to transgene excision. Successful ReMET in F2 progeny is indicated by the loss of yellow body color and both fluorescent marker sequences (Fig. 1A). A silent mutation (TGG > TcG) was engineered into the right homology arm flanking the yReMET constructs so that a single nucleotide change would be left behind following SSA repair, providing evidence that the transgene had in fact been present prior to removal, which would otherwise be indistinguishable from wild-type sequences (Fig. 1A and S1A). To our knowledge, the range of direct repeat substrate lengths over which there is a linear relationship with SSA frequency has not been determined in *D. melanogaster.* Therefore, yReMET lines with either 30, 250, or 500 bp of homologous flanking repeat sequences were created.

**Figure 1.**
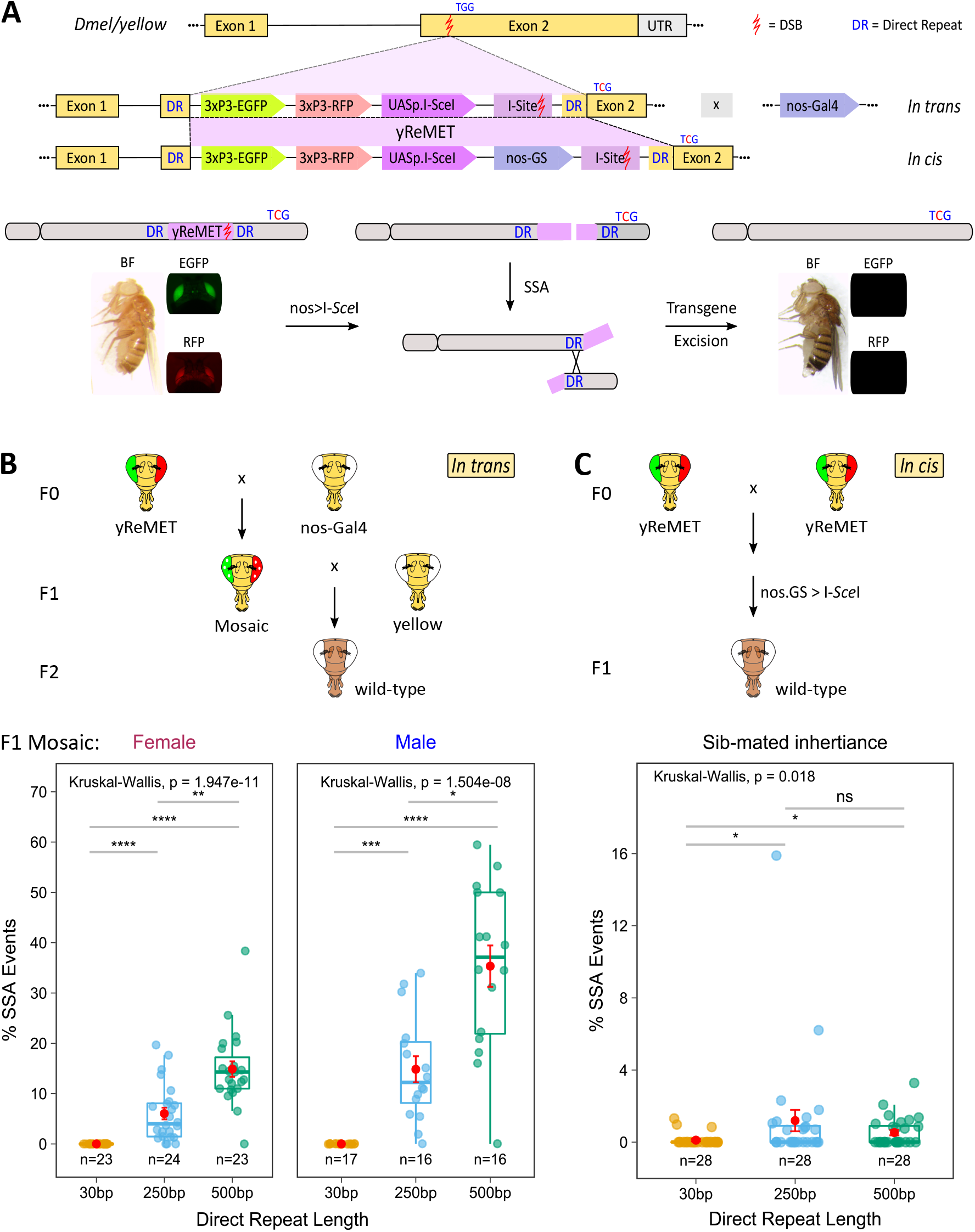
<u>Re</u>peat <u>M</u>ediated <u>E</u>xcision of a <u>T</u>ransgene (ReMET). (**A**) Schematic depicting ReMET constructs (*in trans* and *in cis*) and their removal by SSA. The yReMET construct (shaded in purple) contains EGFP and RFP under the control of independent 3xP3 promoters, and an endonuclease, I-*Sce*I, under the control of UASp. The construct is flanked by a pair of direct repeats (DR) that are 30, 250, or 500 bp in length. Insertion of the ReMET construct into exon 2 of the *yellow* gene creates a null allele that produces yellow body pigmentation with both green and red eye fluorescence. I-*Sce*I may be expressed by crossing with a nos-Gal4 driver line (*in trans*), or by expression from a RU486-inducible nos-GS (*in cis*). Expression of I-*Sce*I generates a DSB at the recognition site (I-Site) located between the DRs, excising the transgene and one of the direct repeats through SSA. SSA-mediated removal of the transgene restores wild type body pigmentation (brown body) with simultaneous loss of eye-specific fluorescence. The engineered synonymous mutation TGG->TcG provides molecular confirmation of ReMET. (**B**) Schematic depicting *in trans* mating scheme and phenotypic identification of germline SSA events. Percentage of F2 progeny exhibiting SSA-mediated transgene excision by DR length, confirmed through wild type body color and the presence of the engineered silent mutation (TcG). Each dot represents a separate pair-mated cross (mean and ± s.e.m. are shown in red), with the sex of the F1 yReMET fly shown above. (**C)** Schematic depicting *in cis* mating scheme and phenotypic identification of germline SSA events. Percentage of F1 progeny exhibiting SSA-mediated transgene excision by DR length, confirmed through wild type body color and the presence of the engineered silent mutation (TcG). Each dot represents a separate pair-mated cross (mean and ± s.e.m. are shown in red). UTR = Untranslated Region; DSB = Double Strand Break; UASp = Upstream Activation Sequence with pTransposase promoter; EGFP = Enhanced Green Fluorescent Protein; RFP = Red Fluorescent Protein; SSA = Single Strand Annealing; p values = * < 0.5, ** < 0.01, *** < 0.001, and **** < 0.0001; ns = not significant.

It is well known that the GeneSwitch system exhibits some leakiness (Scialo et al., 2016). Therefore, the stability of reporter lines containing direct repeats was initially investigated with ReMET constructs lacking the inducible *nos-GS/*GAL4 expression system. Importantly, we did not observe evidence of unintended recombination events after crossing these lines with wild-type flies (Fig. S1B). However, pair-mated reciprocal crosses of F1 flies with a *nos-Gal4* driver line resulted in successful ReMET events (i.e, excision of the transgene mediated by SSA). The frequency of these events was consistent with a linear relationship between SSA and the direct repeat lengths flanking the transgene, with lower rates of excision observed for the 250 bp length and higher rates observed with the 500 bp length (Fig. 1B). We did not observe SSA-mediated excision of the transgene with the 30 bp direct repeat length (Fig. 1B), suggesting that this length may be at or below a minimum effective processing segment (MEPS) length (Shen and Huang, 1986). Interestingly, SSA appeared to occur at a relatively higher mean rate in the male germline, with successful ReMET observed in ∼15% of F2 progeny when the 250 bp direct repeats were present, and in ∼35% of the F2 offspring when the 500 bp direct repeats were present. However, successful ReMET was observed in only ∼6% of the F2 offspring of yReMET females for the 250 direct repeat length, and in ∼15% of progeny when the 500 bp direct repeats were present (Fig. 1B).

As expected, crossing siblings with the complete ReMET construct, which included *nos-GS/*GAL4, in the absence of RU486 resulted in low levels of progeny (0.2 to 0.6%) that exhibited evidence of SSA (Fig. S1C). However, a more surprising result was that significant increases in the frequencies of successful ReMET were not observed in the presence of RU486 (Fig. S1C). ReMET was observed in ∼1% of the F1 offspring with the 250 direct repeat length, and in ∼0.5% of the progeny when the 500 bp direct repeats were present (Fig. 1C). Similar to our earlier results, there was little evidence of SSA with the 30 bp direct repeat length (Fig. 1C). It is possible that the relatively lower levels of SSA observed in these experiments are somehow related to the continual leaky expression of the endonuclease, I-*Sce*I, from the *nos-GS/*GAL4 system throughout development, rather than introduction through a parental cross, as was the case with the previous *in trans* configuration. Nevertheless, the presence of the silent molecular marker, TcG, engineered into the right homology arm provided sequence confirmation of ReMET in all flies exhibiting evidence of SSA (i.e., the loss of yellow body color as well as both fluorescent marker sequences). In order to confirm activation of the *nos-GS* by mifepristone, males containing the full ReMET construct were crossed with a previously published transgenic line (Rong and Golic, 2000), containing a *white* (*w^+^*) reporter gene with an adjacent I-Site sequence, in the presence of increasing concentrations of RU486. As expected, cutting at the site adjacent to the *w^+^* reporter by I-*Sce*I, expressed from the ReMET cassette, deleted all or part of the *w^+^* gene resulting in eye-color mosaicism (Fig. S1D). While I-*Sce*I was expressed at low levels from 5xUAS, even in the absence of treatment with mifepristone, increasing levels of mosaicism were observed in the eyes of F1 progeny depending on the concentration of RU486 present (Fig. S1D). This confirms that the *nos-GS* system present in the full ReMET constructs is functioning properly. Thus, leaky expression of I-*Sce*I from the *nos-GS* system appears to induce SSA at a maximum level, explaining why additional activation with RU486 does not increase ReMET frequencies in the F1 progeny (Fig. S1C). Nevertheless, these experiments demonstrate co-option of the SSA repair pathway for the precise removal of a transgene from *D. melanogaster*.

### Super-Mendelian inheritance of a CRISPR/Cas9-based autonomous homing gene drive

In order to extend our findings demonstrating removal of a transgene to an active gene drive, we reconstructed the previously described CRISPR/Cas9-based yMCR gene drive (Gantz and Bier, 2015; Xu et al., 2020). This gene drive system is based on homology-dependent integration into the X-linked recessive *yellow* locus (Fig. S2A). As per Mendel’s first law, a transgene segregates independently during gamete formation, so that each gamete carries only a single transgenic allele (Fig 2B). However, in genetic drive, allelic conversion transforms heterozygous recessive loss-of-function mutations into homozygous null mutations in the homogametic maternal germline (Fig. 2C), resulting in super-Mendelian inheritance of the drive transgene at frequencies > 50% (Champer et al., 2017; Gantz and Bier, 2015; Xu et al., 2020). This process, which is mediated through repair of the DSB by HR, has also been referred to as a mutagenic chain reaction (MCR). However, other genetic outcomes are also possible (Fig. 2C). Repair of the DSB generated by the Cas9 RNP by the more error prone non-homologous end joining (NHEJ) pathway may sometimes generate resistant alleles. Male flies inheriting resistant alleles may sometimes exhibit a wild-type phenotype, if the mutation does not shift the open reading frame, or a yellow body phenotype, if an out-of-frame mutation occurs (Fig. 2C). Female progeny inheriting these alleles generally do not appear yellow, even if the mutation is out-of-frame, as the paternal wild-type allele is dominant (Fig. 2C). However, female progeny inheriting a maternal allele containing either the transgene or an out-of-frame mutation may occasionally exhibit a yellow phenotype, or mosaic patches of wild-type pigmentation on a largely yellow body. This likely occurs through the persistence of maternally deposited Cas9 RNP in the zygote, which repeatedly cleaves the paternal wild-type allele until it is converted into a mutant allele by NHEJ, resulting in the observed zygotic allelic conversion, or somatic mosaicism (Fig. 2C). While in-frame mutations may sometimes result in a wild-type phenotype, out-of-frame alleles generally produce female progeny with the yellow body pigmentation (Fig. 2C). This non-Mendelian exception resulting from the perdurance of maternally deposited Cas9 RNP has been termed “dominant maternal effect” (Lin and Potter, 2016), and has been observed in gene drive experiments across species (Champer et al., 2018, 2017; Gantz and Bier, 2015; Grunwald et al., 2019; Guichard et al., 2019; Hammond et al., 2016; Xu et al., 2020).

**Figure 2.**
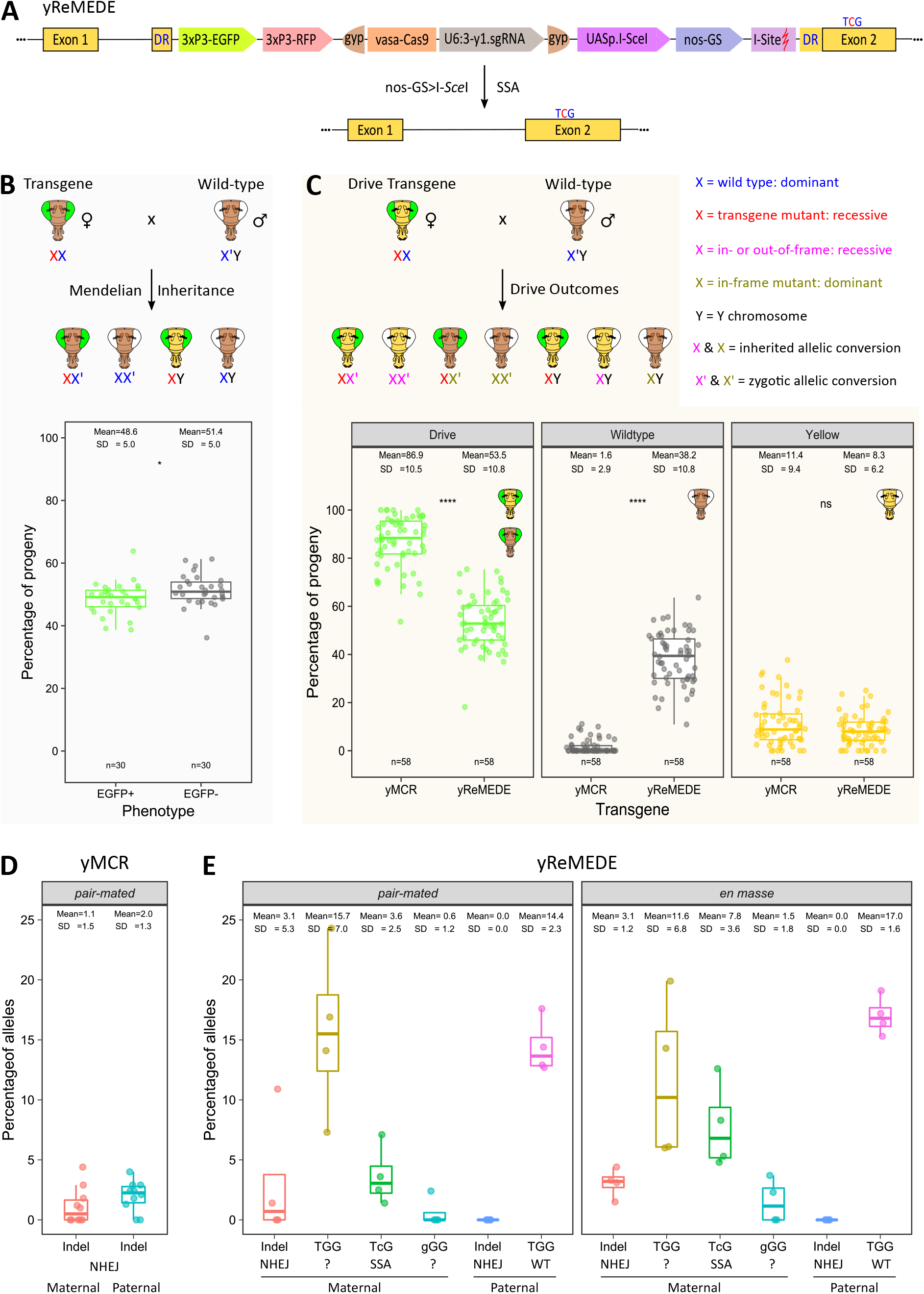
<u>Re</u>peat <u>M</u>ediated <u>E</u>xcision of a <u>D</u>rive <u>E</u>lement (ReMEDE). (**A**) Schematic depicting ReMEDE construct and its removal by SSA. The construct contains components of both the ReMET (*in cis*) and yMCR constructs, including those essential for genetic drive, i.e., Cas9 (under the control of a vasa promoter) and an sgRNA targeting exon 2 of the wild type *yellow* gene (under the control of a U6:3 promoter). The core gene drive components are flanked by a pair of gypsy insulator sequences (gyp). Expression of I-*Sce*I from nos-GS generates a DSB at the I-Site located between the DR, resulting in excision of the transgene by SSA, which is confirmed by the presence of an engineered silent mutation, TcG. (**B**) Schematic depicting maternal inheritance of an X-linked transgene. Mendelian inheritance of the recessive X-linked *yellow* mutant allele (generated through transgene insertion) from a heterozygous mother produces wild type body pigmentation in heterozygous female progeny, but a yellow phenotype in the hemizygous male progeny. Maternal inheritance of a transgene expressing GFP inserted into the yMCR and yReMEDE sgRNA target site. The percentage of progeny exhibiting GFP+ or GFP-phenotypes in replicate pair-mated crosses of heterozygous F1 females with wild type males are shown. Each dot represents a separate pair-mated cross. (**C)** When the transgene is a gene drive, multiple Mendelian outcomes, as well as non-Mendelian exceptions, are possible. When the heterozygous drive allele present in the F1 female creates a DSB in its wild type counterpart, HR-mediated repair results in allelic conversion increasing transgene frequency and generating super-Mendelian patterns of inheritance (green eyes). However, DSBs repaired by the more erroneous NHEJ may generate either recessive out-of-frame null alleles, or dominant in-frame alleles. Non-Mendelian exceptions may also occur due to the zygotic perdurance of maternally deposited Cas9 RNPs generating DSBs in the paternal wild type alleles, resulting in either in-frame or out-of-frame repair, which generates the additional possibilities illustrated. Maternal inheritance of either the yMCR or yReMEDE alleles. The percentage of progeny exhibiting drive, wild type, or yellow phenotypes in replicate pair-mated crosses of heterozygous F1 females with wild type males are shown. Each dot represents a separate pair-mated cross (**D**) Sequencing results for the *yellow* gene in wild type progeny of pair-mated crosses involving F1 yMCR females and wild type males, sorted by parental lineage. (**E**) Sequencing results for the *yellow* gene in wild type progeny of pair-mated or *en masse* crosses involving F1 yReMEDE females and wild type males, sorted by parental lineage. The sequence present at the PAM site is shown, as well as the DNA repair pathway presumed to be responsible for its generation. NHEJ = Non-Homologous End Joining; SSA = Single Strand Annealing; p values = ** < 0.01, *** < 0.001, and **** < 0.0001; ns = not significant.

Our MCR drive was modified to include *3xP3-EGFP* (MCR-EGFP) so that we could track the appearance of resistant alleles, both functional and non-functional (Fig. S2A). To validate the drive, we performed a series of independent single pair-mated outcrosses (n=58) with heterozygous F1 females and wild-type males. The F1 females were generated by crossing yMCR males to wild-type females. The mean transmission rate of the drive allele to the F2 progeny over all crosses was ∼87% (Fig. 2C). The mean rate of offspring exhibiting a yellow mutant phenotype was ∼11%, with the remaining ∼2% of all F2 progeny displaying a wild-type phenotype (Fig. 2C). It should be noted that these percentages are consistent with previously reported yMCR inheritance patterns in *D. melanogaster* (Champer et al., 2018, 2017; Gantz and Bier, 2015; Xu et al., 2020).

### <u>Re</u>peat <u>M</u>ediated <u>E</u>xcision of a <u>D</u>rive <u>E</u>lement (ReMEDE)

The yMCR construct was next recombined into the yReMET (250 bp) reporter line by standard methods to generate a yReMEDE line (Fig. 2A and Fig. S2B). Based on results regarding direct repeat substrate length and SSA frequency (see above), all subsequent experiments employed a direct repeat length of 250 bp. F1 females were generated by crossing yReMEDE males to wild-type females. The performance of the yReMEDE system was then assessed in a parallel series of outcrosses (n = 58), where heterozygous F1 yReMEDE females were pair-mated with wild-type males. The allelic inheritance patterns of the resulting progeny were then compared to those resulting from the outcrosses with the F1 yMCR females (Fig. 2C). Both the genotypes and phenotypes of progeny that are expected to result from crosses involving F1 yReMEDE females are shown in Figure S3. Successful ReMEDE was indicated by loss of fluorescent marker genes and reversion to wild-type body pigmentation in the F2 progeny (Fig. S3). As expected, the number of individuals with wild-type phenotypes increased significantly, but somewhat unexpectedly these comprised ∼38% of the total yReMEDE progeny over all crosses (Fig. 2C). Further, this significant increase in wild-type progeny correlated with a significant decrease in the overall mean transmission rate of the drive allele (∼54%; Fig. 2C and S2C). While we considered the hypothesis that incorporation of the ReMEDE components unintentionally disabled Cas9 resulting in uncut alleles, this was inconsistent with multiple observations. First, in single pair mated crosses (n = 30) of flies containing a transgene lacking drive elements, but inserted into the same location of the *yellow* gene, more than half of the F2 flies inherited the transgene at rates of less than 50%, with only a single replicate nominally exceeding the 60% threshold (Fig. 2B and Data S1). In contrast, transmission of the yReMEDE allele ranged from 18% to 75%, with rates over 60% being observed in 15 separate replicate crosses (Fig. 2C and Data S1). Further, the overall mean inheritance of the transgene was significantly lower than that of the ReMEDE construct (p value = 0.01; Fig. 2B and C). More importantly, yellow F1 females were generated by crossing yReMEDE males with wild-type females (Fig. 2C), which could not occur in absence of Cas9 mediated cutting. Additionally, the overall percentage of progeny exhibiting resistant alleles (i.e., yellow body color) in the yReMEDE F2 generations did not significantly differ from those observed in the yMCR F2 flies (Fig. 2C and Data S1). If the Cas9 were no longer active, the transgenic construct would then be subject to normal Mendelian inheritance, with only GFP positive yellow males generated from the crosses (Fig. 2B). Thus, a more plausible hypothesis consistent with these results is that both ReMEDE and allelic conversion are occurring simultaneously.

Phenotypic scoring of the F2 yReMEDE flies serves as a proxy for the approximate rate of SSA. This is particularly true with regard to the observed increases in numbers of male wild-type progeny generated through outcrosses of F1 yReMEDE females. In males, X-linked inheritance is exclusively maternal, necessarily suggesting germline conversion mediated by SSA (Fig. S2C and S3). Nonetheless, sequencing over the Cas9 target site in all wild-type individuals resulting from multiple pair-mated outcrosses of the F1 yReMEDE females, for the silent mutation (TGG > TcG) engineered into the right homology arm of the construct, confirmed transgene (i.e., the drive element) excision by SSA. By design, this mutation also functions as an engineered resistant allele in the critical protospacer adjacent motif (PAM), essential for Cas9-mediated cleavage (Fig. 2A). The altered PAM sequence prevents self-targeting that might lead to the formation of undesirable NHEJ-mediated resistant alleles or allelic conversion within the ReMEDE allele. Although pair mated crosses with F1 yMCR females rarely produced wild-type phenotypes (Fig. 2C and Data S1), sequencing confirmed them to be indels, generated either by NHEJ acting on a maternally inherited allele, or a dominant maternal effect acting on a paternally inherited allele (Fig. 2D). Overall, these indel sequences occurred at a frequency of ∼1-2% in the inherited alleles (Fig. 2D and Data S1). Alleles containing indels, generated either through NHEJ or dominant maternal effect, occurred with similar frequency (∼3%) in the wild-type progeny of F1 yReMEDE females (Fig. 2E and Data S1). However, sequencing also confirmed SSA-mediated events in ∼4% of the maternally inherited wild-type alleles in the F2 yReMEDE flies (Fig. 2E and Data S1), which were not present in F2 yMCR flies. Paternally inherited wild-type alleles also increased to ∼14% in F2 yReMEDE flies (Fig. 2E and Data S1). However, in contrast with paternally inherited alleles in the wild-type F2 yMCR flies, which contained only indels (Fig. 2D and Data S1), paternal alleles in the wild-type offspring of yReMEDE flies remained intact or uncut, which is consistent with limited or no maternal deposition of Cas9 RNP (Fig. 2E and Data S1).

We were only able to confirm ∼4% of the wild-type alleles resulted from ReMEDE. These alleles were necessarily contributed by the F1 maternal germline (i.e., present in F2 male progeny), and carried the TcG marker sequence (Fig. 2E, S3 and Data S1). However, wild-type alleles containing a wild-type TGG or gGG sequence (possibly produced through erroneous repair) were also present (Fig. 2E, S3 and Data S1). It should be noted that F2 female progeny also inherited alleles with the TcG marker, but ascertaining the precise origin of these alleles is more difficult (Fig. S3). Thus, to investigate the origin of alleles containing the TcG sequence, as well as the TGG and gGG sequences, further we sequenced all progeny generated from a single pair-mated cross with two sets of primers (Pair-mated cross #4; Data S1). The first primer set encompassed the PAM sequence, while the second set encompassed the entire transgene, with binding sites located outside of the construct’s left and right homology arms (Fig. S2A and S2B). The use of both primer sets permitted us to ascertain the zygosity of both drive (ReMEDE) and non-drive alleles (wild-type or containing indels) present in the homogametic female progeny. The results confirmed that only ∼3% of the wild-type alleles in heterozygous females carried the engineered TcG marker, with a larger ∼17% exhibiting the original wild-type TGG sequence (Pair-mated cross #4, Data S1). In contrast, all ReMEDE drive alleles, present in either male or female progeny, were positive for the purposely engineered TcG variant of the PAM sequence (Pair-mated cross #4, Data S1).

Replicate *en masse* crosses involving multiple F1 yReMEDE females and wild-type males yielded an average transmission rate of ∼53% for the drive allele, with ∼42% of the progeny exhibiting a wild-type phenotype (Fig. S2D), numbers very similar to those observed in pair-mated crosses (Fig. 2C). Similar percentages were observed in a series of additional replicate crosses performed in the presence of RU486 (Fig. S2D). Sequencing over the Cas9 target site in wild-type individuals for the silent mutation (TGG > TcG), engineered into the right homology arm of the construct, again provided confirmation of drive element excision by SSA (Fig. 2E and Data S1). Taken together, these experiments demonstrate proof-of-principle for successful co-option of the highly conserved SSA repair pathway for the precise autocatalytic excision of an active gene drive.

### ReMEDE converts the MCR-EGFP gene drive to wild-type alleles in population cage studies

In order to determine if the ReMEDE system would convert the gene drive element into wild-type alleles over multiple generations, we conducted cage trials with either the yReMEDE or yMCR drive flies. The cage trials were performed in accordance with those described in previously published studies (Bishop et al., 2022; Xu et al., 2020). We evenly divided six hundred male and female virgins into three replicate cages (200/cage). Within each cage, 25% of the flies were wild-type and 75% homozygous for either yMCR or yReMEDE. At each generation (n), approximately 150 flies/cage were randomly selected for scoring (i.e., fluorescent markers and body color), and another randomly selected 150 flies/cage seeded into the subsequent generation (n + 1) of cages. The frequency of flies with unmodified yMCR alleles (EGFP+) decreased only modestly, still representing a significant portion of the population after 14 generations (Fig. 3A). In contrast, the frequency of flies exhibiting yReMEDE drive phenotypes (EGFP+, RFP+, and yellow body color) decreased gradually, and were completely eliminated in 10-14 generations (Fig. 3B). Reciprocally, the frequency of flies exhibiting a wild-type phenotype increased gradually, representing 100% of the population after 10-14 generations (Fig. 3B). These experiments suggest that all gene drive elements carrying ReMEDE will eventually be converted to wild-type alleles given enough generations.

**Figure 3.**
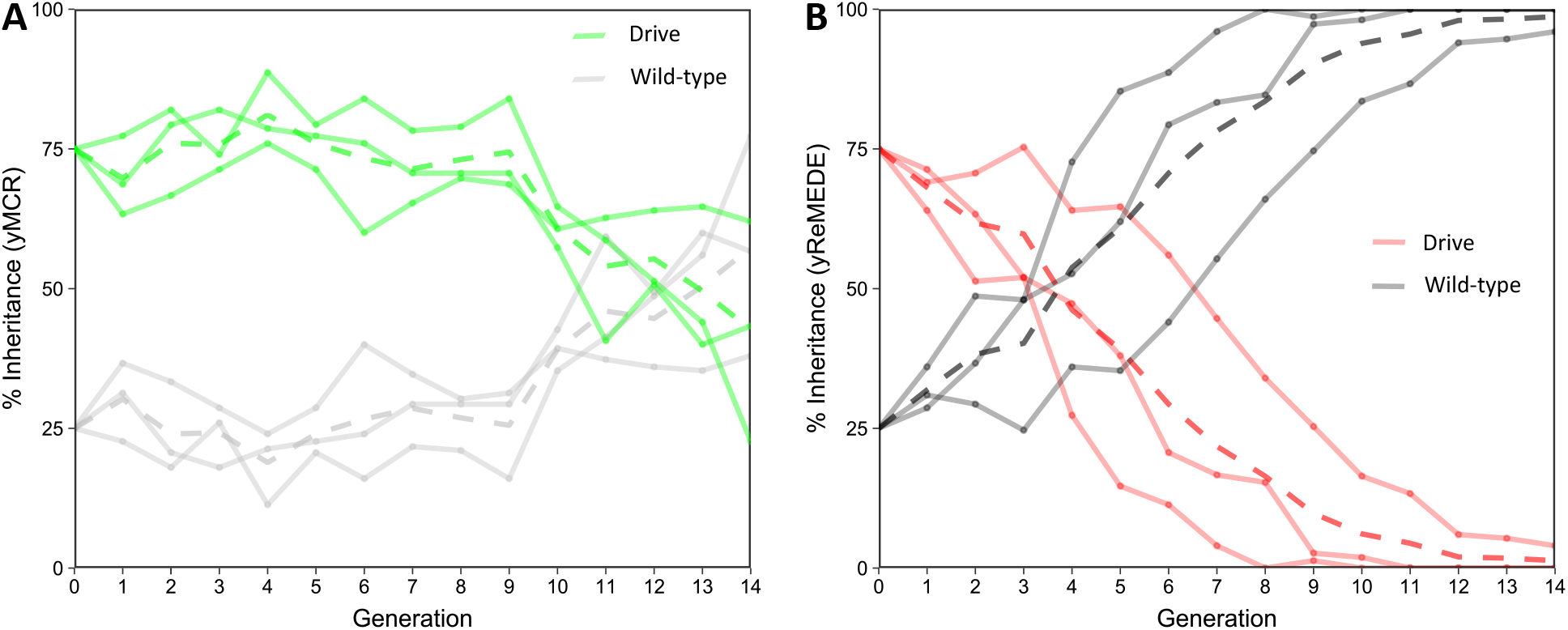
Population cage trials with either yMCR or yReMEDE gene drives. Graphs show the prevalence of drive and wild type phenotypes (exhibited by both male and female flies) over time in cage trials involving either (**A)** yMCR (green lines) or (**B**) yReMEDE (red lines). Solid lines represent single replicate cages, while dotted lines represent mean values.

### ReMEDE-mediated conversion of a threshold-independent gene drive to wild-type

The conversion of the yMCR gene drive element into wild-type alleles by ReMEDE over multiple generations suggests that the system will perform similarly in converting a low-threshold gene drive into wild-type alleles. The *yellow* gene target site of the yMCR gene drive renders this element threshold dependent. First, the target cut site is susceptible to erroneous repair by NHEJ or other error-prone DNA repair pathways, which may disrupt the PAM sequence, generating alleles that can no longer be targeted by the Cas9 ribonucleoprotein (RNP), i.e., resistant alleles. Second, mutations in the *yellow* gene adversely affect male mating success (Barker, 1962; Bastock, 1956; Drapeau et al., 2006; Massey et al., 2019; Sturtevant, 1915; Xu et al., 2020). In order to generate mathematical models to predict the dynamics of a low or no-threshold drive containing ReMEDE over multiple generations, we determined the mating cost associated with *yellow* mutant flies, containing an insertion at the same site targeted by the drive allele. *D. melanogaster* engages in a series of courtship rituals (e.g., tapping, chasing, genital licking, singing, etc.) prior to successful copulation. A previous study has shown that the lower mating success of *yellow* males in comparison to wild-type males does not result from any observable difference in the amount of time spent courting, but rather from delayed initiation of copulation (Massey et al., 2019). However, these experiments were conducted with Canton-S wild-type flies. In our population studies, it was necessary to use the *w^1118^*strain as wild type, in order to score eye-specific fluorescence. The *white* (*w*) gene has also been associated with copulation success, and *w^1118^*flies display reduced courtship activity (Xiao et al., 2017). Thus, we calculated courtship indices (the proportion of time that a male was engaged in courtship activities divided by the total duration of observation) for wild-type (*w^1118^*) and mutant (*w^1118^;yellow*) flies. Although wild-type males displayed reduced courting activities with wild-type females, the courtship of mutant *yellow* females was even more markedly reduced (Fig. S5A). Male *yellow* mutants had no success mating with wild-type females in these trials; thus, a courtship index was not recorded for these crosses (Fig. S5A). However, the courtship index of mutant *yellow* males was approximately 0.5, when calculated from crosses with mutant *yellow* females. Given the results of the single pair courtship assays, mating costs associated with mutant *yellow* males were instead calculated from competitive assays, where a wild-type virgin female was provided with both a wild-type male and mutant *yellow* male. In the 30 crosses performed for these assays, a single *yellow* male, exhibiting a courtship index of ∼0.5, successfully copulated (Fig. S5A). Based on the results of these experiments, we estimated the mating cost associated with mutant *yellow* males to be 29/30 or 0.97 (Fig. S5A).

We have previously generated mathematical models for SSA-based excision of transgenes (Chae et al., 2022; Zapletal et al., 2021). These initial parameters were used to simulate allele frequencies for both autosomal and X-linked inheritance patterns (Fig. S5C and Supplementary Methods), and then modified to include empirical values derived from crosses involving yMCR (δ = NHEJ and mating cost of 0.97 for yellow males) and yReMEDE (α = SSA, γ = SSA resistance, and ε = engineered allele persistence) flies. This model was then used to run simulations of various scenarios. In conjunction with these simulations, we conducted multi-generation population cage trials to assess the impact of mating costs and resistant alleles associated with the *yellow* gene target site of the yMCR gene drive. Replicate cages were seeded with flies, such that yMCR females or males comprised 10% of the population, with the other 90% being virgin wild-type males or females; a level that might be consistent with the introduction of a low-threshold drive element. The scoring of each generation (n) and seeding of subsequent generations (n + 1) was performed as described above. In each of the cages, individuals exhibiting fluorescence and yellow body phenotypes were rapidly lost from the population, regardless of sex (Fig. S5B). Simulations conducted with similar parameters matched the observed performance of the drive element (g) in population cage studies (Fig. S5D). In an attempt to remove the mating costs associated with the *yellow* gene target site, additional cages were seeded with flies in which yMCR females or males again comprised 10% of the population, but with the other 90% being males or females homozygous for the null *yellow* allele (i.e., a mutant population). The frequency of the EGFP+ cassette (i.e., the yMCR drive allele) increased in the population, ultimately achieving 15-60% penetrance when introduced by female flies, and 30-80% penetrance when introduced by the male flies (Fig. S5B). Simulations of male releases derived with similar parameters roughly approximated the trends observed in the population cage trials (Fig. 4A).

**Figure 4.**
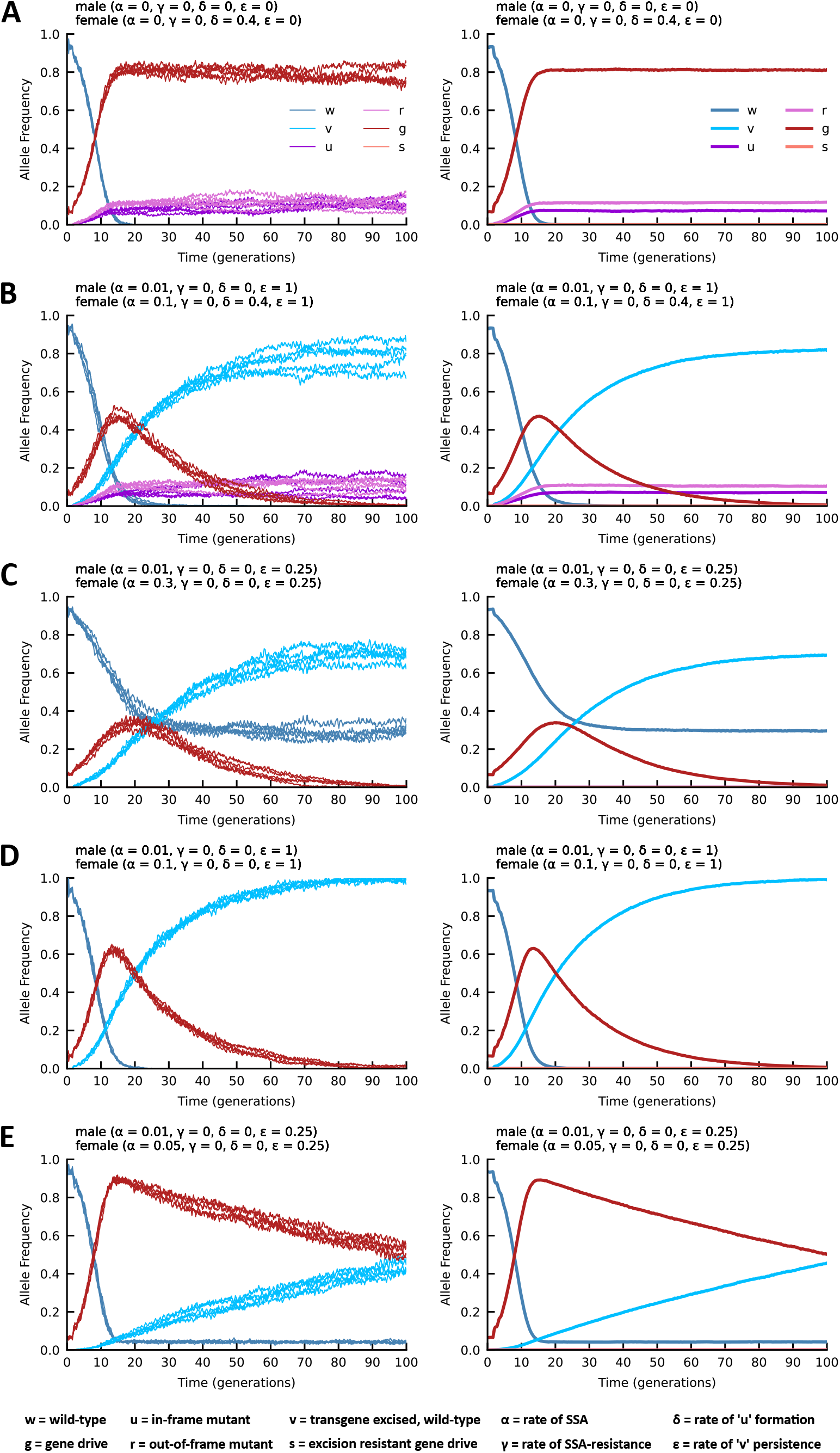
Stochastic modeling of ReMEDE technology performance in a population. All models were run with the X-linked inheritance module that outputs allele frequencies in two plots. One plot shows five simulations randomly selected from 100 independently run simulations (Left), while the other plot shows the mean of all simulations (Right). All simulations were run in a low threshold release scenario (10% of flies containing a drive allele) and in the absence of mating costs. (**A**) Simulation of yMCR gene drive, generating resistant alleles at frequencies comparable to those observed experimentally, but in the absence of reduced courtship activity and mating success. (**B**) Simulation of yReMEDE where SSA converts approximately 10% of the drive alleles to wild type. The generation of resistant alleles and other parameters occur at frequencies comparable to those shown in A. (**C**) Simulation of a gene drive containing ReMEDE that generates no resistance alleles, but with SSA converting approximately 30% of the drive alleles to wild type; a quarter of which retain the engineered resistant allele in the PAM site (ε = 0.25). (**D**) Simulation of a gene drive containing ReMEDE that generates no resistance alleles, but with SSA converting approximately 10% of the drive alleles to wild type, all of which retain the engineered resistant allele in the PAM site (ε = 1) (**E**) Simulation of a gene drive containing ReMEDE that generates no resistance alleles, but with SSA converting a modest ∼5% of the drive alleles to wild type, a quarter of which retain the engineered resistant allele in the PAM site (ε = 0.25).

Additional mathematical modeling removing all mating costs associated the yReMEDE drive, and with SSA (female α = 0.1, male α = 0.01) occurring at ∼10%, where all events are marked (TcG) by the engineered resistant allele (v; ε = 1), results in a maximum ∼50% drive allele frequency after about 15 generations, which are then progressively replaced by drive-resistant TcG-marked wild-type alleles (v) (Fig. 4B). However, several studies have analyzed and overcome the challenge of resistant alleles, either by targeting essential genes, using multiple guide RNAs, recoding target genes, or by incorporating a combination of these strategies to ensure that only homozygous drive or wild-type allele combinations remain viable (Champer et al., 2020, 2018; Kandul et al., 2021; Kyrou et al., 2018). In lieu of these empirical developments, we simulated yReMEDE population dynamics in the absence of both mating costs and resistant alleles (δ = 0), with three different SSA frequencies. If approximately 30% of maternally inherited alleles are formerly drive alleles that were converted to wild-type alleles by SSA, of which 25% retain the engineered TcG resistant allele (v; ε = 0.25), w alleles (native wild type) and v alleles (drive-resistant wild type) persist in the population, while the drive alleles (g) are progressively eliminated (Fig. 4C). Alternatively, when SSA occurs at ∼10%, with all events retaining the TcG marker (ε = 1), allelic conversions of wild-type alleles (w) by drive alleles achieve a maximum frequency of ∼60% in < 15 generations, but are then subsequently entirely replaced by drive-resistant alleles (v) via SSA (Fig. 4D). When SSA occurs at the minimal rate of ∼5%, with only 25% of these retaining the TcG marker (ε = 0.25), the frequency of the drive allele (g) quickly reaches ∼90% in < 15 generations, but is subsequently replaced by engineered drive-resistant alleles (v) after several more generations (Fig. 4E). Therefore, taken together, stochastic modeling informed by empirical data, indicates that a mere 5-10% of SSA events where only one-fourth of the events remain drive-resistant (v), allow for efficient propagation of a low-threshold drive with eventual replacement by engineered drive-resistant wild-type alleles.

## Discussion

Early successes mitigating the negative effects of resistance alleles on drive efficiency (Carballar-Lejarazú et al., 2020; Hammond et al., 2021, 2017; Kyrou et al., 2018; Unckless et al., 2017) suggest that the development of technologies for limiting the spread of drives through a population may be necessary, if such tools are ever to be successfully tested and deployed in the field. Indeed, several technological solutions have recently been proposed for homing drives, with the design and demonstration of various braking systems and confinable gene drive configurations representing a particularly positive development in this regard (Basgall et al., 2018; Chae et al., 2020; DiCarlo et al., 2015; Esvelt et al., 2014; Gantz and Bier, 2016; Taxiarchi et al., 2021; Wu et al., 2016). Nonetheless, it remains unclear if any of the currently proposed technologies will ultimately prove to be an adequate solution. First, many of these strategies are premised on the introduction of additional independent transgenic genetic elements. These elements share a number of similarities with gene drives themselves, and as a result their efficacy may be influenced by the same parameters as those affecting the homing drives they are designed to stop (i.e, gene conversion efficiency, fitness costs, timing, etc.), and indeed this has been observed in several laboratory trials and simulations (Alphey and Bonsall, 2014; Deredec et al., 2008; Tanaka et al., 2017; Unckless et al., 2015; Xu et al., 2020). Second, the introduction of additional transgenic elements may complicate risk assessment, as it might not be possible to extrapolate with a high enough degree of confidence how an autonomous drive might function in a particular ecological environment based on field testing in a split configuration, or in the presence of an independent genetic braking element. Third, no strategy satisfactorily addresses the critical requirement for removal of transgenes from natural populations, should that become necessary. Rather, the removal of multiple independent transgenes at the conclusion of a field trial, in any sort of acceptable time frame, would likely require inundation through the sustained release of wild-type individuals. We, therefore, sought to develop a solution where the excision of the transgene would be autocatalytic over the period of genetic drive, eventually restoring the wild-type population. By design, drive activity is meant to be gradually reduced as drive alleles are converted to wild-type alleles. We have previously proposed co-opting the SSA repair mechanism for this purpose (Zapletal et al., 2021, Chae et al., 2022).

Here, we demonstrate proof-of-concept using a self-eliminating version of the yMCR drive element (Gantz and Bier, 2015) targeting the *yellow* gene of *D. melanogaster*. The system, which we termed ReMEDE, significantly increased wild-type progeny. Crossing F1 yMCR females with wild-type males rarely produced wild-type individuals (Fig. 2C and Data S1). Sequencing these individuals confirmed them to be the result of maternally inherited indels generated through NHEJ, or a dominant maternal effect, acting on the paternally inherited allele (Fig. 2D). Overall, these indel sequences were present in ∼1-2% of the inherited alleles (Fig. 2D and Data S1). These results are consistent with previous studies with the yMCR drive where the occurrence of in-frame, functional, mutations were also found to be relatively rare (Champer et al., 2017; Gantz and Bier, 2015; Xu et al., 2020). Alleles with indels generated either through NHEJ or dominant maternal effect were found to occur with similar frequency (∼3%) in the progeny of F1 yReMEDE females (Fig. 2E and Data S1). However, in addition to these, sequencing confirmed SSA-mediated events in ∼4% of the maternally inherited alleles (Fig. 2E and Data S1), confirming a genetic basis for observed increases in wild-type progeny in the crosses involving yReMEDE flies. Paternally inherited wild-type alleles also increased to ∼14% in the yReMEDE offspring (Fig. 2E and Data S1). However, in contrast with the wild-type progeny of the yMCR flies, where all paternal alleles had been converted into indels, all of the paternal alleles present in the wild-type progeny of the yReMEDE flies remained intact or uncut, suggesting that the maternal deposition of Cas9 RNP was somehow prevented (Fig. 2E and Data S1). One possible explanation for this is that ReMEDE-mediated removal of the gene drive, suppressed Cas9 RNP perdurance (Fig. 2D, S2C, 2E, and Data S1), which would have otherwise resulted in zygotic allelic conversion.

Sequencing wild-type individuals generated from the pair-mated crosses of yMCR females did not reveal any alleles with a TGG PAM sequence (Fig. 2D and Data S1), suggesting that non-drive alleles are repeatedly cut with the Cas9 nuclease, eventually resulting in either allelic conversion or the formation of a resistant allele. Indels were found with similar frequency in the wild-type offspring of both yReMEDE and yMCR females (Fig. 2D, 2E and Data S1), as were resistant alleles (i.e., yellow progeny) in the F2 generations (Fig. 2C and Data S1), suggesting that the Cas9 remains active in the yReMEDE drive allele. Rather, the primary difference, evident from the results of single pair mated crosses, was the presence of wild-type flies with alleles containing either a TGG or TcG sequence at the PAM site in the F2 yReMEDE generation, which were not found in the F2 yMCR generation (Fig. 2D, 2E and Data S1). Sequencing of intact ReMEDE drive alleles revealed only the purposely engineered TcG variant of the PAM sequence (Pair-mated cross #4, Data S1). Thus, overall, these results are consistent with both drive conversion and SSA happening simultaneously in the ReMEDE flies. While we can only speculate on the origin of the wild-type alleles containing the TGG sequence, one possibility is that these are uncut alleles occurring because of ReMEDE-mediated removal of the drive allele. However, another hypothesis is that ReMEDE occurs with subsequent conversion of TcG to TGG in the F1 heterozygous maternal germline, via synthesis-dependent strand annealing (SDSA) and mismatch repair (MMR; see Fig. S4 for model).

There are two generally accepted models for the repair of DSBs by HR, double strand break repair (DSBR) and synthesis-dependent strand annealing (SDSA). Both models begin with the resection of the 5’ termini flanking the DSB. This generates long single-stranded 3’ tails, one of which invades the homologous duplex template sequence resulting in the formation of a D-loop. After this step, the two models diverge wherein the DSBR model is characterized by formation of double Holliday junctions (dHJs), the resolution of which may result in either crossover or non-crossover products (Szostak et al., 1983). SDSA is mediated by new synthesis, disassociation of the recently extended strand, and annealing with the other 3’ tail in the broken chromosome, exclusively generating non-crossover products (Fig. S4). While the DSBR model appears to be preferred in meiotic cells, evidence increasingly suggests that the SDSA model is the major pathway of DSB repair in mitotic cells (Andersen and Sekelsky, 2010). In some cases, MMR machinery appears to be capable of recognizing and repairing mismatches present in the transient heteroduplex DNA (hetDNA) intermediates formed during the repair process (Jiricny, 2006). However, the newly synthesized strands, produced in both the DSBR and SDSA models, may also form hetDNA that can be detected and repaired through MMR. In either case, loss of heterozygosity would be possible through nonreciprocal transfer of information from the template strand (Hum and Jinks-Robertson, 2017). In the ReMEDE system, DSBs generated by I-*Sce*I occur between engineered direct repeats increasing the likelihood that they will be repaired by SSA, whereas those generated by Cas9 do not occur between direct repeats making it more likely that HR will be chosen. Occurrences of wild-type PAM sequences (TGG) in the yReMEDE lines, but not in yReMET lines that were devoid of Cas9, might suggest that it is the cleavage mediated by Cas9 that is necessary for SDSA, and subsequently MMR, resulting in reversion of the engineered marker TcG to TGG. Sequencing of progeny from single pair-mated crosses revealed that all of the ReMEDE drive alleles present retained the engineered silent mutation, suggesting that these reversions would have to happen following excision of the ReMEDE construct (Data S1).

Our previous work on this topic was entirely conceptual, as such it was impossible to know how this technology would perform in an actual biological organism. However, one concern that we considered a distinct possibility was that SSA might be too efficient at removing the transgene, to the point that gene drive might not occur. In order to address this possibility, we placed the second nuclease (I-SceI) under the control of an inducible gene switch system. However, this system is leaky, and we continue to see expression of I-SceI (Fig. S1D). To our knowledge, all inducible systems exhibit some level of leakiness. In any case, these concerns ultimately proved to be unfounded, as SSA occurred at relatively low levels, even when higher levels of the nuclease were produced (Fig. S1C). These results suggest that there is no need for an inducible system in future iterations of the technology, which in any case would have been difficult to implement in a field setting. In practice, SSA re-constitutes wild-type alleles at relatively low levels, which is desirable as the wild-type population is only gradually restored, permitting a period of genetic drive. In multi-generational population cage studies, the ReMEDE drive element was completely eliminated, and a wild-type population restored in 10 to 14 generations (Fig. 3). A number of factors related to the performance of the yMCR gene drive (i.e., mating defects, resistant allele formation, etc.) prevented us from conducting population-level studies under conditions that would mimic the release of a low-threshold gene drive (Fig. S5). However, simulations with experimental parameters obtained from this study suggest a similar dynamic would occur with low threshold drives. Simulations of yReMEDE population dynamics, in the absence of both mating costs and resistant allele formation, determined that only 5% of SSA-mediated recombination events, even with only a fraction of these retaining the engineered PAM site, would be sufficient to permit the initial propagation of a low threshold drive, followed by its eventual replacement with wild-type alleles (Fig. 4E). These frequencies already fall below those observed in single pair mating experiments, suggesting that the ReMEDE technology may be capable of halting even the most robust gene drive, eventually reverting the population back to a wild-type state. Given the limitations of the yMCR gene drive, future work will focus on employing the ReMEDE system in the context of a more robust low threshold gene drive. However, the present study constitutes an effective proof-of-principle for the ReMEDE concept as a viable technology for controlling autonomous homing gene drives. The ReMEDE system has a number of theoretically desirable features as a technological safeguard for gene drives, including the potential to restore the original wild-type population, and addressing the issue of transgene persistence in natural populations. The ReMEDE concept should be considered along with other strategies currently under investigation for controlling, neutralizing, halting, deleting, etc., gene drives. Particularly, as early modeling efforts suggest that threshold-independent drives, similar to those that have already been developed in the malaria vector *Anopheles gambiae,* may be difficult to control with braking systems (Rode et al., 2020). While other strategies have also been demonstrated in laboratory settings, it is too soon to know if any of these will be successful in field settings (Basgall et al., 2018; Chae et al., 2020; Esvelt et al., 2014; Gantz and Bier, 2016). Further, it is entirely plausible that no mitigation strategy will prove ideal for every possible scenario involving gene drives. Thus, prudence would appear to dictate that multiple gene drive mitigation strategies are pursued further at this juncture.

## Acknowledgements

We thank Dr. Omar Akbari at the University of California San Diego for reviewing the manuscript and providing helpful comments. The graphic cartoon art of the *Drosophila* head used in the figures was downloaded as a vector file from Zenodo (Somers, 2020) and modified.

## Author Contributions

KM, ZA, and PC conceptualized the study; PC and KM designed the research; PC, RM, and KM performed the experiments and analyzed the data; JZ, MN, and PC developed and ran the model simulations; PC and KM wrote the manuscript; KM, PC, and ZA revised the manuscript. All authors approved the final manuscript.

## Data Availability

The data for this study are provided in the supplementary information/source data file provided with this paper.

## Code Availability

All files pertaining to the generation of equations, model execution, and plotting of charts can be found at the following link: https://github.com/mln27/Python-CRISPR1-1-SEM-gene-drive

## Supplementary Methods

### Courtship Assays

A construct containing EGFP under the control of a 3xP3 promoter was inserted into exon 2 of the *D. melanogaster yellow* gene at the drive element (yMCR or yReMEDE) target site, generating a null mutation. The courtship behavior of yellow flies was then assessed in both noncompetitive and competitive mating crosses with wild-type (*w^1118^*) flies conducted in specially designed fly arenas, as described previously (Boutros et al., 2017). Briefly, female and male flies were gently aspirated into the circular arenas of the apparatus. Mating behavior was then captured by video over a five-hour duration. Any copulation between a male and female fly observed during the video analysis was subsequently scored as a successful mating. Courtship studies were conducted at room temperature over a similar time range during the day (12 hour light/dark cycle) with a standard corn-meal/oatmeal based medium.

### Population Studies

All multi-generational cage trials were conducted at 25 C (12 hour light/dark cycle) in standard 250 mL rearing bottles containing a standard corn-meal/oatmeal based medium. For each genotype, bottles were seeded with equal numbers of unmated male and female flies. After 5 days, this initial generation (n) of flies was removed from the bottles. Subsequent generations (n+1) were then collected and randomly divided into two equal pools. One pool was seeded into a fresh cage (n+2), while the other was scored and further analyzed.

### Modeling

Mathematical modeling was performed with methods that have been described previously (Chae et al., 2022), but with the following modifications. In this study, the continuous addition of offspring into the adult mating pool was replaced with distinct generations. A modified version of the widely used lumped age-class model of mosquito life-history (Hancock and Godfray, 2007; Deredec et al., 2011; Sánchez C. et al., 2020) was incorporated to more fully account for the fly’s four different states: egg, larvae, pupae, and adult (male and female). In order to adapt the model to *Drosophila,* the following parameters were assigned to the fly’s life-history stages. Egg production per day (day[-1]) = 100; duration of egg stage (days) =1; duration of larval stage (days) = 4; duration of pupa stage (days) = 5 (Fernández-Moreno et al., 2007; Ong et al., 2015; Mołoń et al., 2020); daily mortality risk of egg stage (day[-1]) = 0.15; daily mortality risk of pupa stage (day[-1]) = 0.15 (Durdevic et al., 2018; Dylan Shropshire et al., 2018; Layton et al., 2019); daily mortality risk of adult stage (day[-1]) = 0.263; and daily population growth rate (day[-1]) = 1.196 (Horváth et al., 2016). We also developed a stochastic version of the previously described methods where genotype-specific daily egg production follows a multivariate Poisson distribution informed by parental genotypes and the “Equation Generation” module. Survival and death events follow binomial distributions at the population level.

#### Equation Generation

Equations for determining the number of offspring with each genotype were derived as described previously (Chae et al., 2022), but with the following modifications. An additional factor of epsilon (ε) was incorporated into the equations to capture the rate of ‘v’ persistence. Where ‘v’ represents a wild-type allele resulting from the excision of a previously inserted drive element, but engineered to contain a Cas9 target site that is resistant to re-cutting. The formula 1-ε was then used to represent a ‘v’ allele that has been converted into the original wild-type allele (w) sequence by mismatch repair. In order to simulate the X-linked inheritance of the ReMEDE drive construct, the probability equations for the heterogametic males were simplified to include a single allele. All possible combinations of the two female alleles and one male allele were diagramed and probabilities computed for each.

#### Model Accessibility

Details regarding the execution of modeling methods and interpretation of results have been provided previously (Chae et al., 2022). All files pertaining to the generation of equations, model execution, and plotting of charts can be found at the following link: https://github.com/mln27/Python-CRISPR1-1-SEM-gene-drive

## Supplementary Figure Legends

**Figure S1.**
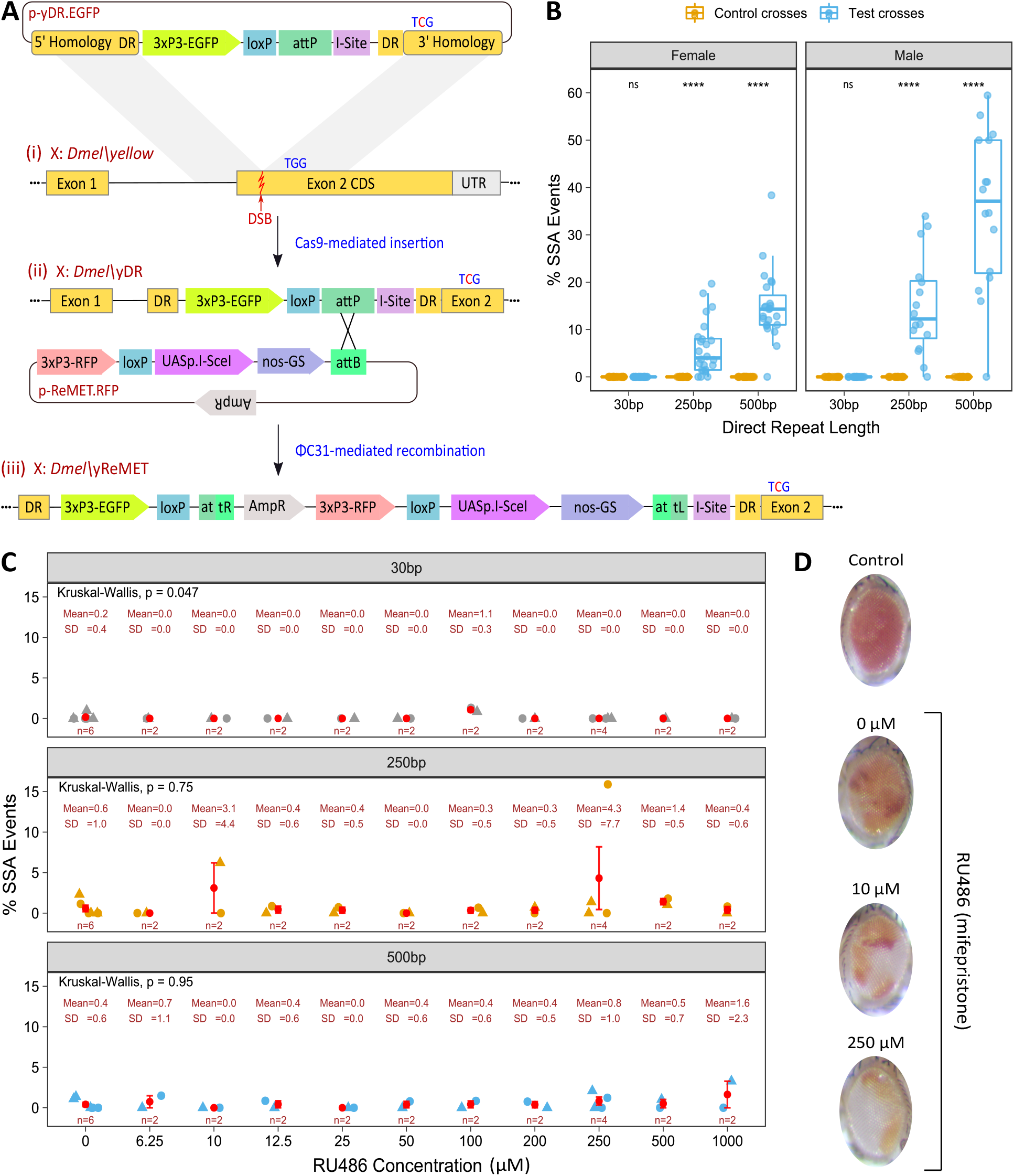
Generation of yReMET lines, transgene excision, and characterization of RU486 inducible promoter. (**A**) Schematic depicting the two-step process for the generation of yReMET (*in cis*) transgenic lines. In the first step, a donor plasmid, p-yDR.EGFP, with DRs of varying lengths (30, 250 or 500 bp), identical in sequence to the 3’ end of the 5’ homology arm, were inserted into the *Drosophila melanogaster yellow* gene at a Cas9 target site. The donor construct contained 3xP3-EGFP, which served as a marker of transgenesis, a recognition site for the I-*Sce*I endonuclease (I-Site), a loxP site, and an attP landing site. The resulting yDR transgenic fly lines were also engineered to contain the silent PAM mutation TGG->TcG. In the second step, a donor plasmid, p-ReMET.RFP, containing an attB site was inserted into the yDR lines through ϕC31-mediated recombination. The p-ReMET.RFP donor contained 3xP3-RFP, a loxP site, as well as the nos-GS and UASp.I-*Sce*I components. In another configuration, used for the *in trans* experiments, only the UASp.I-*Sce*I is included and not the nos-GS. Integration of the p-ReMET.RFP plasmid into the yDR lines generated the various yReMET lines containing DRs of different lengths. (**B**) Percentage of F2 progeny exhibiting SSA-mediated transgene excision by DR length and sex, confirmed through the presence of wild type body pigmentation and the engineered silent mutation (TcG). Gold dots represent individual pair-mated crosses of F1 yReMET (*in trans* configuration) flies with wild type flies. Blue dots represent individual pair-mated crosses of F1 yReMET (*in trans* configuration) flies with transgenic nos-Gal4 flies. P-value **** < 0.0001; ns = not significant. (**C**) Percentage of F1 progeny exhibiting SSA-mediated transgene excision by DR length and RU486 concentration, confirmed through the presence of wild type body pigmentation and the engineered silent mutation (TcG). Each dot/triangle represents a separate pair-mated cross (mean and ± s.e.m. are shown in red). (**D**) Images of adult eye pigmentation in F1 progeny resulting from pair-mated crosses of yReMET (*in cis* configuration) flies with flies containing a *white* (*w^+^*) reporter gene with an adjacent I-Site sequence. Images are representative of the average level of mosaicism observed for the given concentration of RU486.

**Figure S2.**
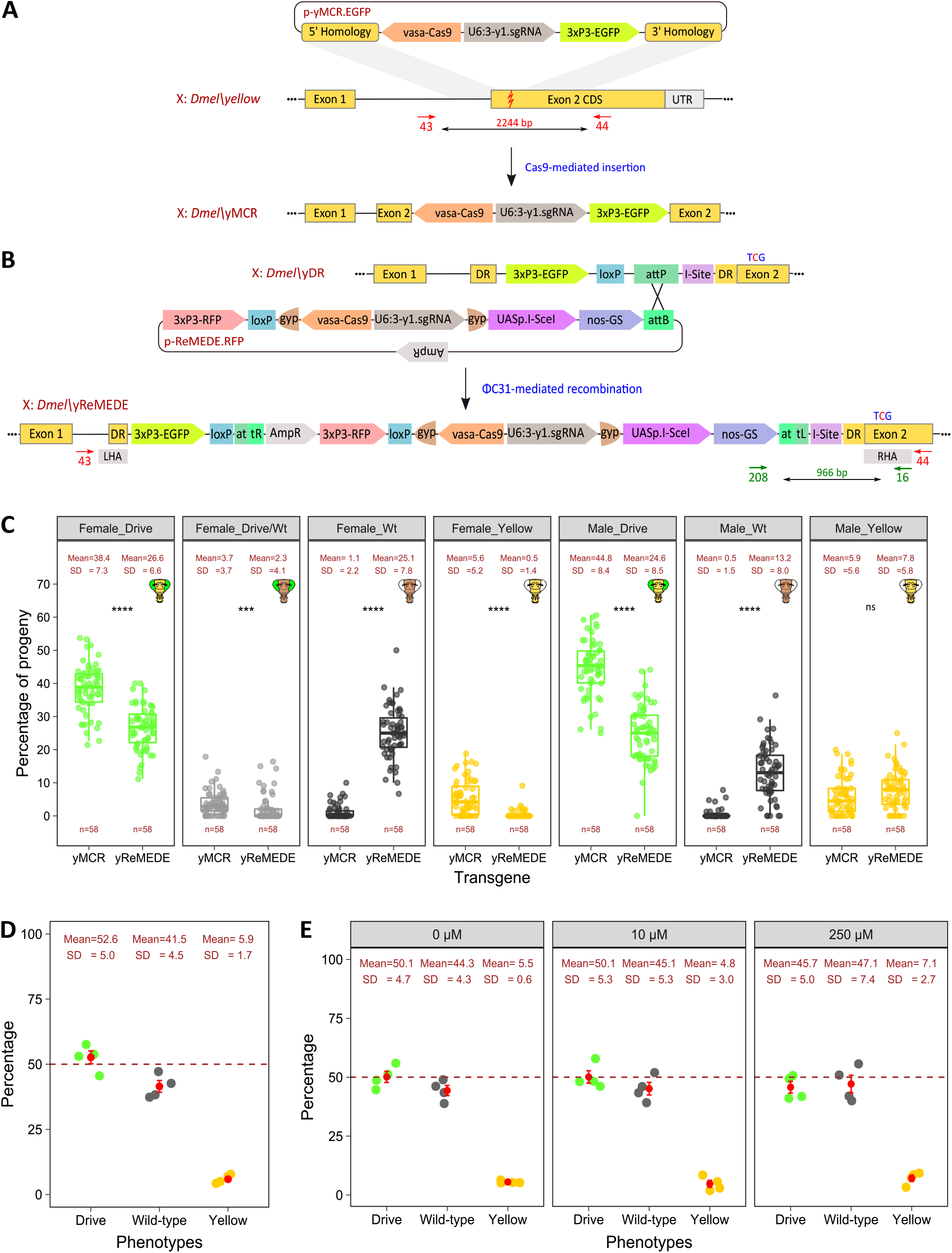
Generation and characterization of yReMEDE line. (**A**) Schematic depicting insertion of p-yMCR.EGFP into the *yellow* gene of *Drosophila melanogaster*. The plasmid p-yMCR.EGFP contains Cas9 (under the control of a vasa promoter), an sgRNA targeting exon 2 of the wild type *yellow* gene (under the control of a U6:3 promoter), and EGFP (under the control of the eye-specific 3xP3 promoter). The primer annealing sites 43 and 44 are located outside the homology arms in order to permit genotyping of target alleles in flies exhibiting a wild type phenotype. (**B**) Schematic depicting the generation of the yReMEDE line through ϕC31-mediated recombination of p-ReMEDE.RFP into the yDR line. The plasmid p-ReMEDE.RFP contains the yMCR drive element flanked by a pair of gypsy insulators (gyp), nos-GS > UASp-I-*Sce*I, a loxP site, and 3xP3-RFP. Primer annealing sites 208 and 16 permit genotyping of the PAM site in the drive allele. (**C**) Maternal and paternal inheritance of drive elements. Percentage of progeny exhibiting scored phenotypes (i.e., body pigmentation and fluorescent eye color) as indicated by cartoon fly heads. Each dot represents a separate pair-mated cross. (**D**) Percentage of progeny exhibiting wild type phenotypes in replicate *en masse* crosses of yReMEDE flies. Each dot represents a separate *en masse* cross (mean and ± s.e.m. are shown in red). (**E**) Percentage of progeny exhibiting wild type phenotypes in replicate *en masse* crosses of yReMEDE flies in the presence of RU486 at the concentrations specified. LHA = Left Homology Arm; RHA = Right Homology Arm; p values = ** < 0.01, *** < 0.001, and **** < 0.0001; ns = not significant.

**Figure S3.**
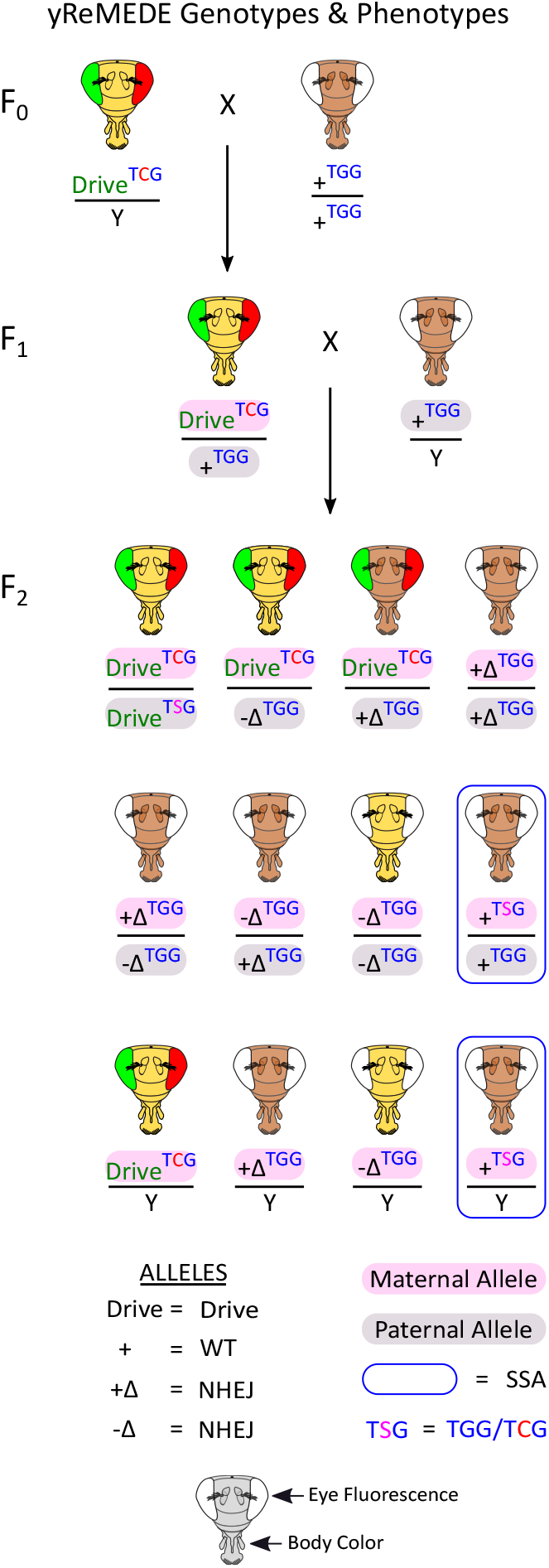
Matrilineal inheritance of wild type alleles in progeny of females carrying yReMEDE. Mating scheme for generation of F1 yReMEDE females and their F2 progeny. Images of fly heads indicate scored phenotypes of eye fluorescence (white = no marker, green = EGFP, red = RFP) and body color (brown = + or +Δ, yellow = Drive or -Δ). Possible allelic combinations are shown below head images (ReMEDE = Drive, wild type = +, in-frame indel = +Δ, out-of-frame indel = -Δ). Male flies are indicated by the presence of the Y chromosome. Blue boxes enclose genotypes and phenotypes that confirm SSA-mediated transgene excision, regardless of the PAM sequence.

**Figure S4.**
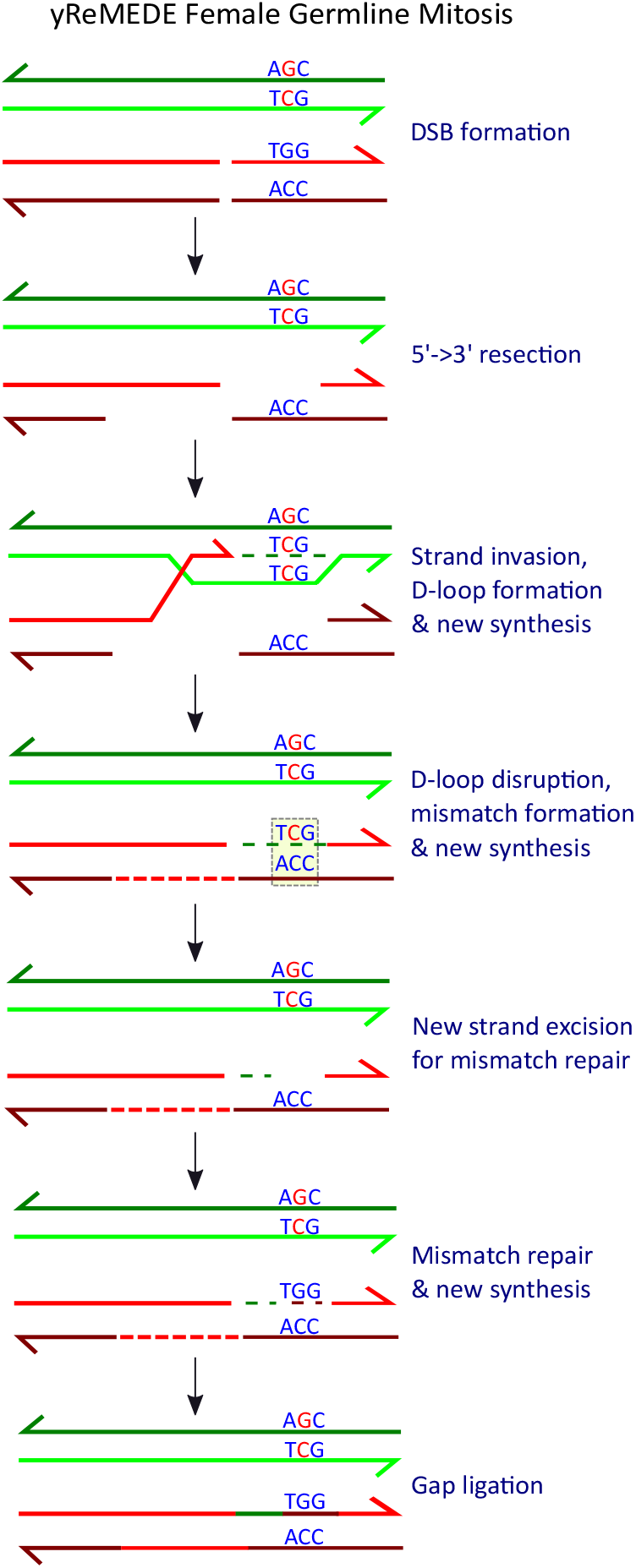
Model for reversion of the engineered TcG PAM sequence to wild-type. The SDSA model appears to be the primary DSB repair mechanism in mitotic cells and is shown for simplicity. Production of Cas9 RNP from the gene drive (donor) allele (shown in green) generates a DSB in the wild type (recipient) allele (shown in red). Resection of the 5’ DNA ends produces 3’ single-stranded tails. Strand invasion of the donor drive allele by one of the recipient allele’s 3’ tail results in base pairing of complementary strands generating a tract of heteroduplex DNA (hetDNA). This process displaces the originally duplexed strand forming a displacement (D)-loop. Elongation of the invading 3’ end increases the size of the D-loop, eventually producing a sequence that extends beyond the DSB. Dismantling of the D-loop frees the newly synthesized sequence to anneal with the other 3’ tail of the recipient allele, generating a new tract of hetDNA with the opposite strand, i.e., SDSA. Mismatches present in the hetDNA are repaired by MMR. Excision followed by resynthesis and ligation repairs the mis-paired bases, restoring the original wild type sequence of the recipient and eliminating heterozygosity at the locus. SDSA = Synthesis Mediated Strand Annealing, DSB = Double Strand Break, RNP = RiboNucleoProtein.

**Figure S5.**
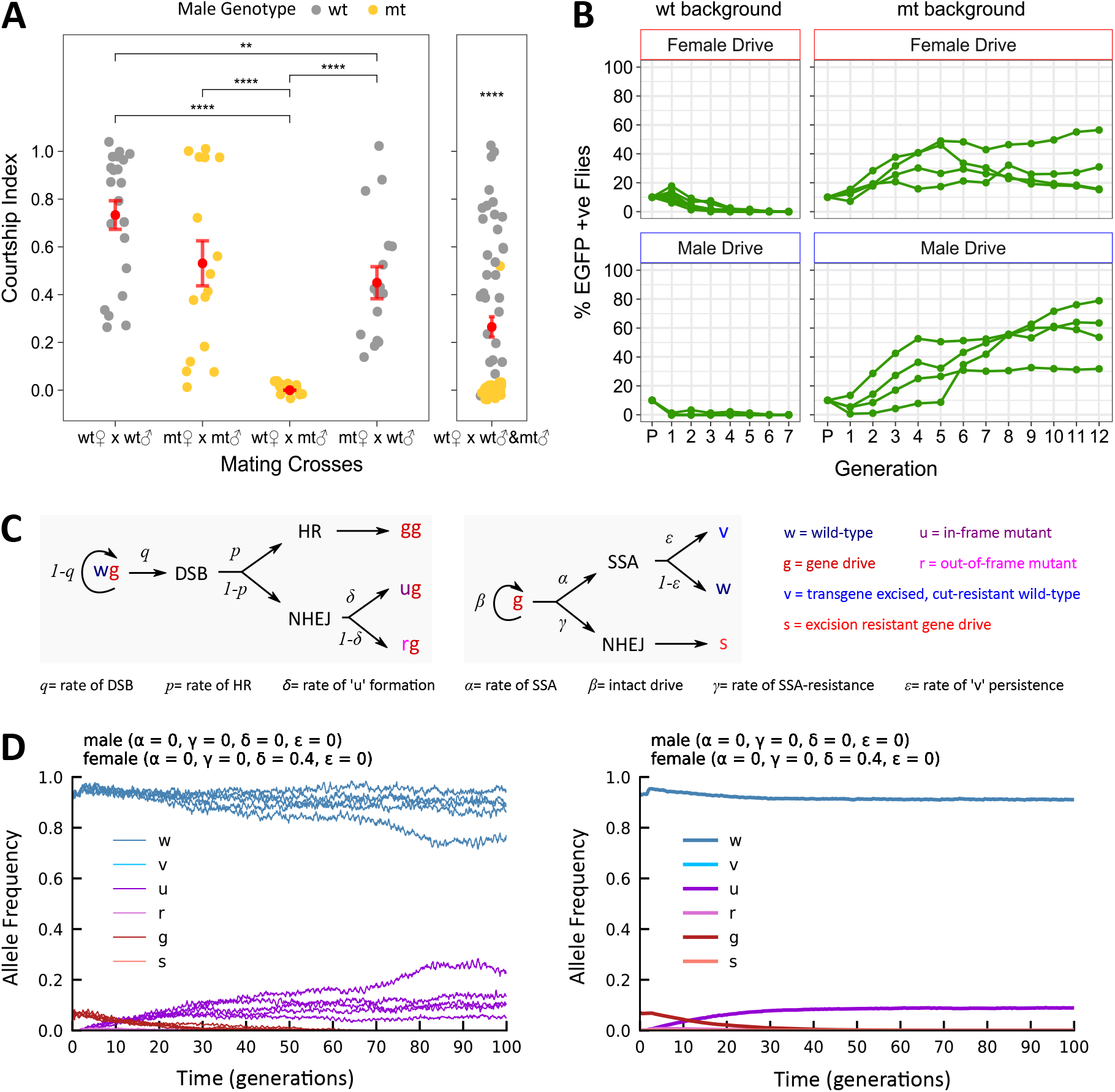
Performance of the yMCR gene drive in a low threshold release scenario. (**A**) Courtship indices calculated for *yellow* mutant (mt) and *w^1118^* (wt) flies in the crossing schemes shown. P-values = ** < 0.01, and **** < 0.0001. (**B**) Cage trials with the yMCR gene drive in either a *w^1118^* (wt) or *yellow* mutant (mt) population. F1 females or males carrying the yMCR allele initially made up 10% of the total population. Graphs show the prevalence of the drive phenotype (green line) in both male and female progeny over time. Each line represents a separate replicate cage trial. (**C**) Theoretical model of allelic conversion dynamics occurring within the F1 female germline during oogenesis. In gene drive, a DSB occurs independently at each susceptible target allele (w) with a probability of *q*, after which repair occurs by HR with a probability of *p,* or by NHEJ with a probability of *1-p*. HR results in allelic conversion of w to g, while NHEJ produces in-frame alleles (u) with a probability of δ, or out-of-frame alleles (r) with a probability of 1-δ. In the presence of ReMEDE, a second DSB occurs and is repaired by SSA with a probability of α, or by NHEJ with a probability of γ. SSA retains the engineered silent PAM mutation (TcG) with a probability of ε, or TcG reverts back to TGG through MMR with a probability of 1-ε. (**D**) Stochastic modeling of the yMCR gene drive in a wild type population. All models were run with the X-linked inheritance module that outputs allele frequencies in two plots. One plot shows five simulations randomly selected from 100 independently run simulations (Left), while the other plot shows the mean of all simulations (Right). Simulations of the yMCR gene drive were run in a low threshold release scenario (10% of flies containing a drive allele), and generating resistant alleles at frequencies (δ = 0.4) comparable to those observed experimentally. Similarly, mating cost parameters were set at levels consistent with empirically determined values for yellow male (0.97) and female (0.31) flies in a wild type population.

**S6.**
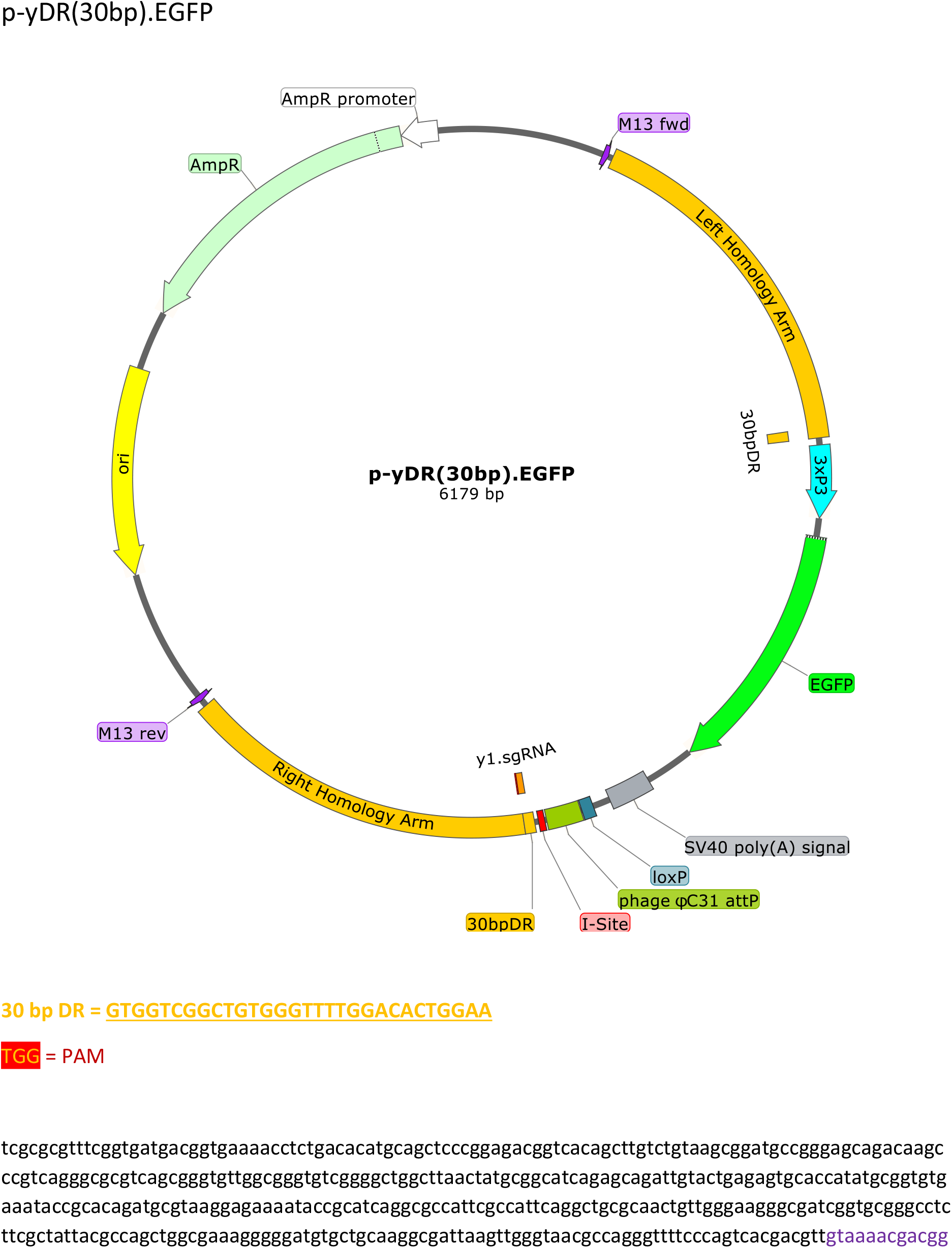
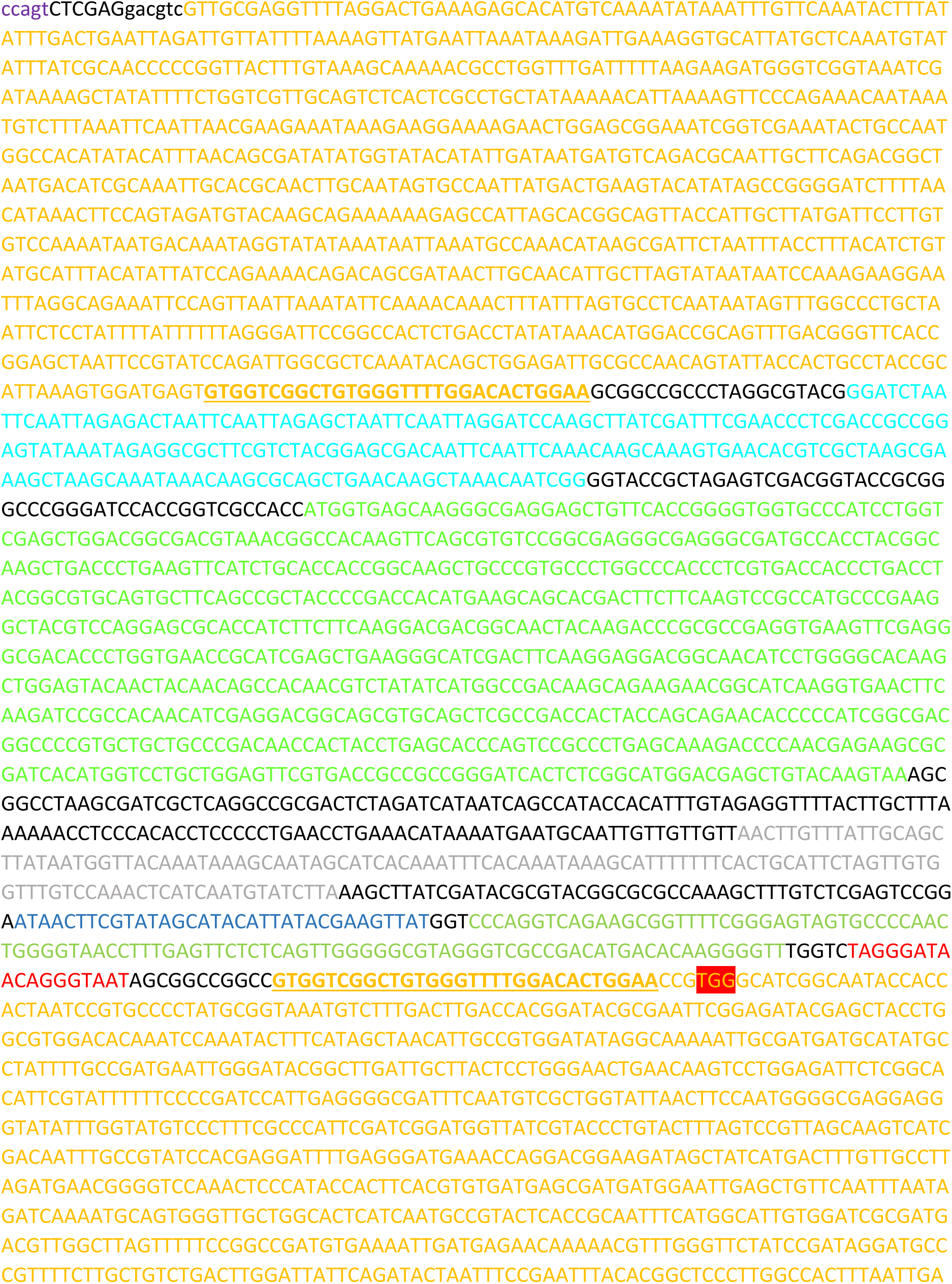
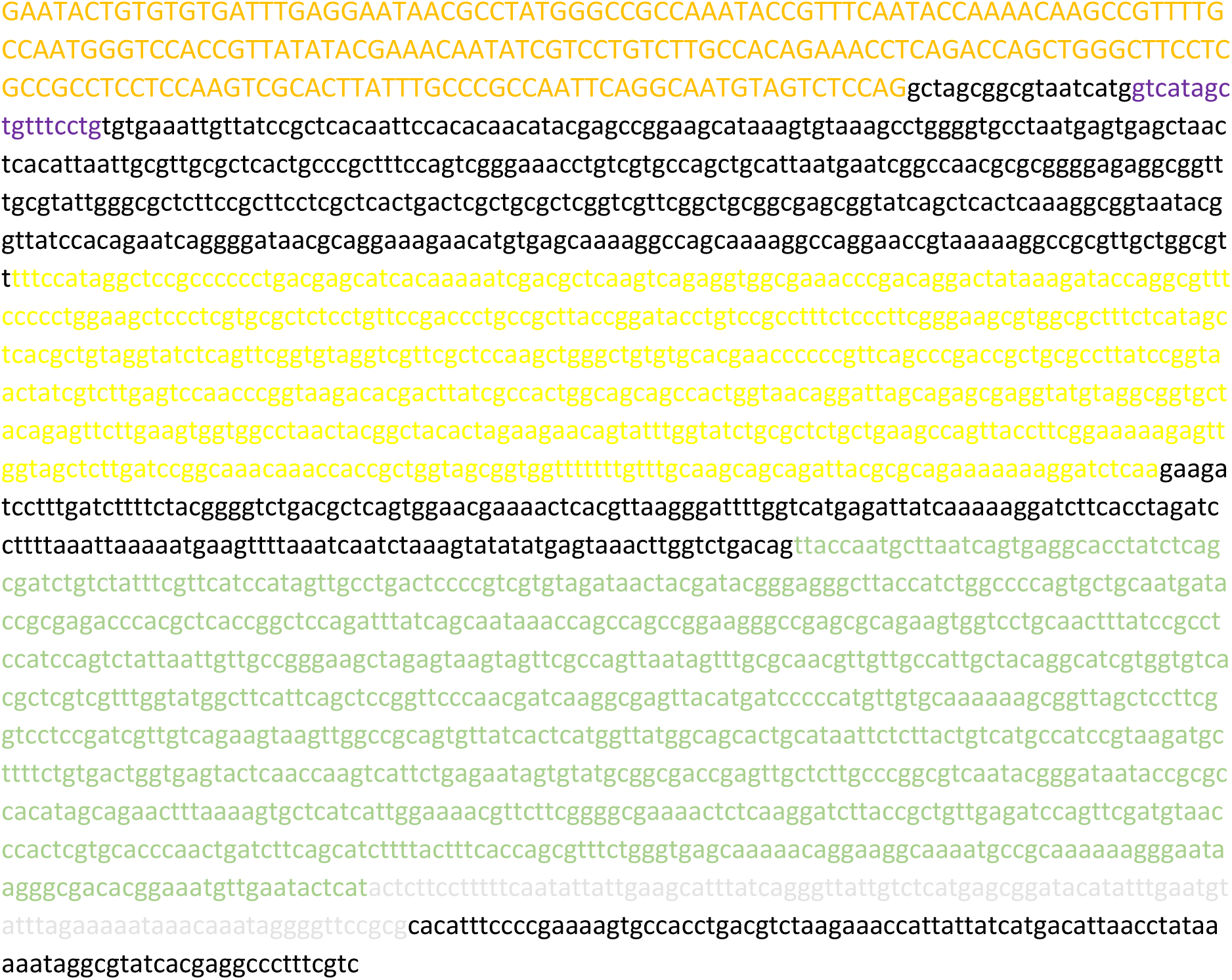
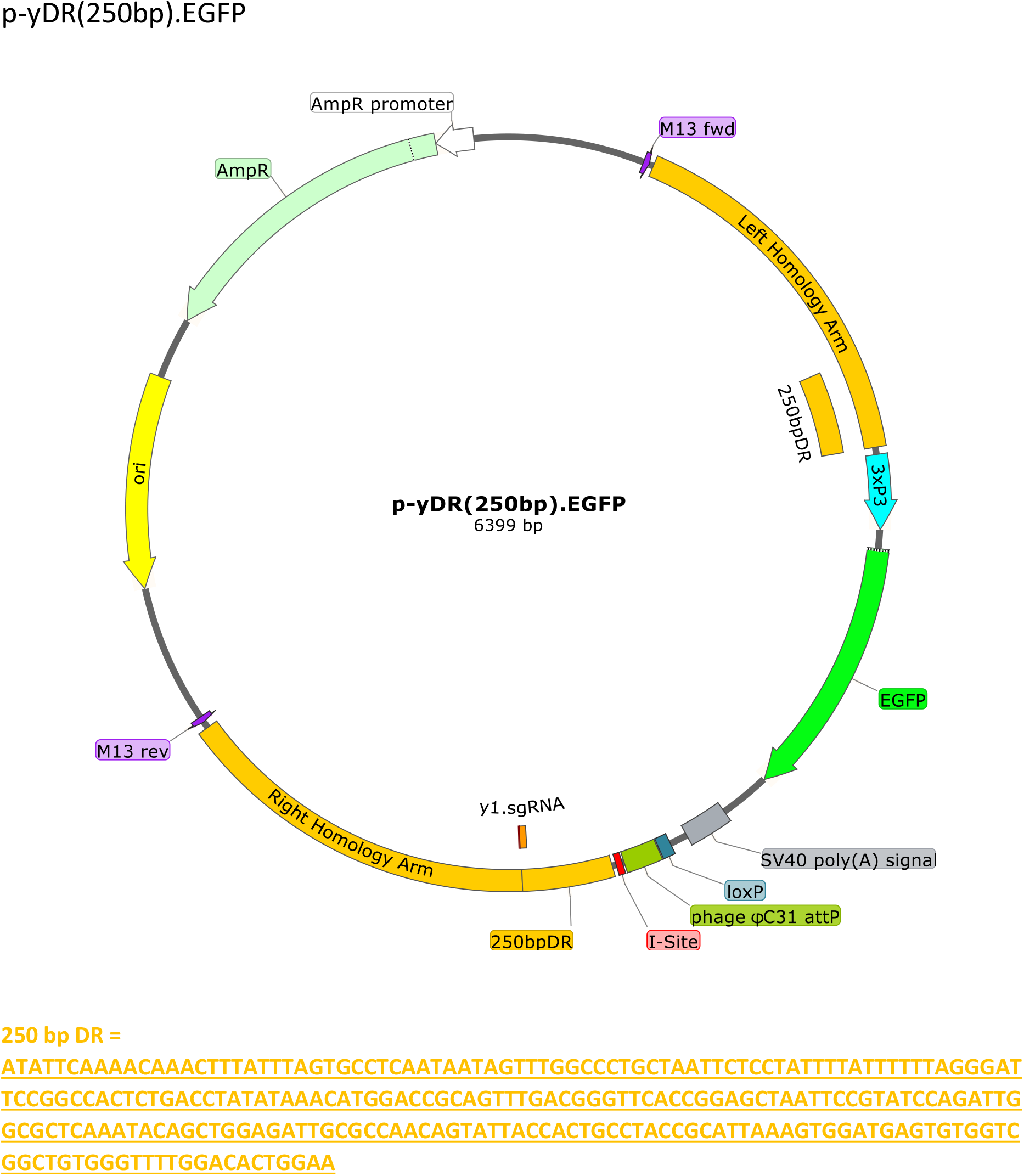
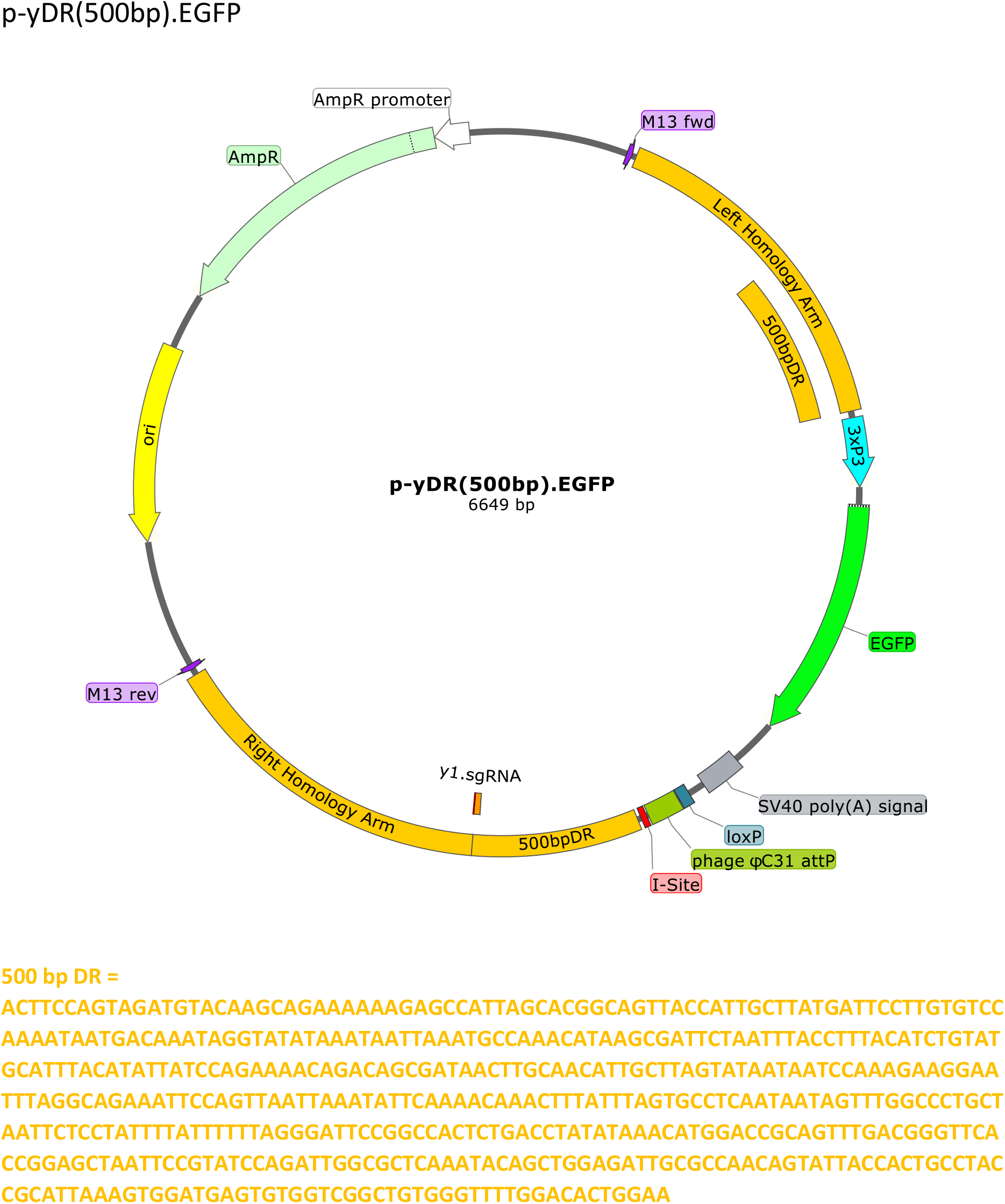
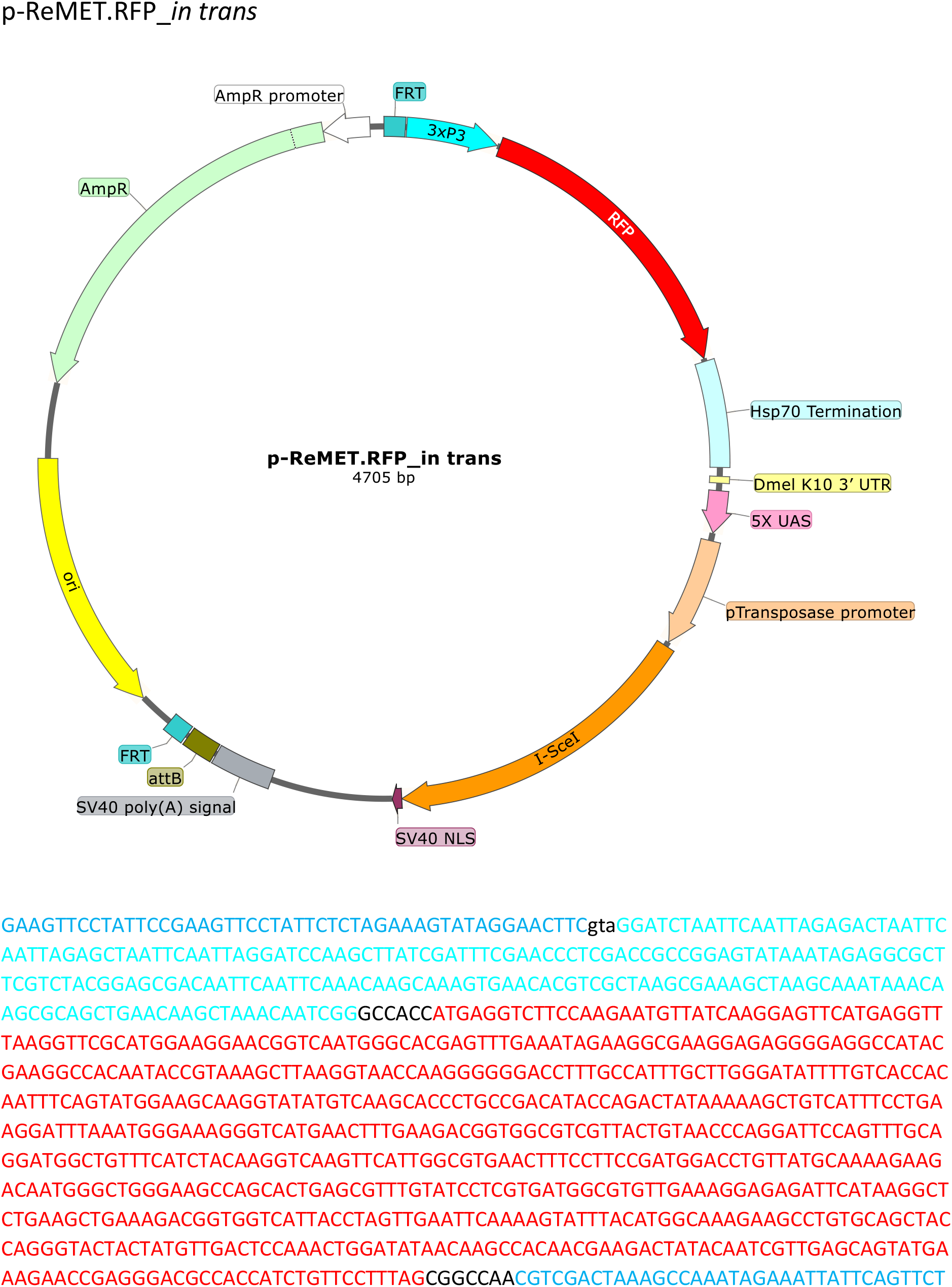
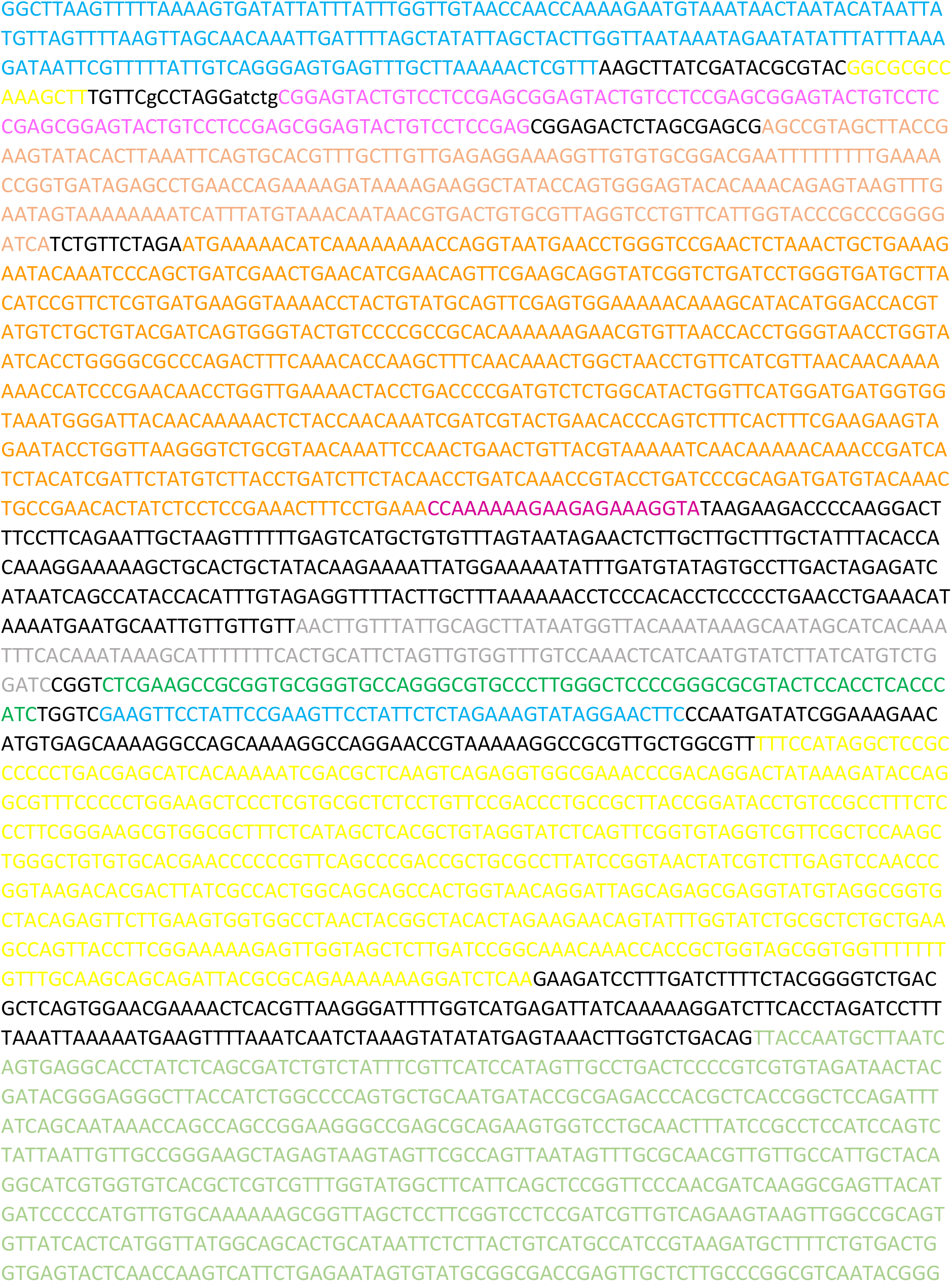
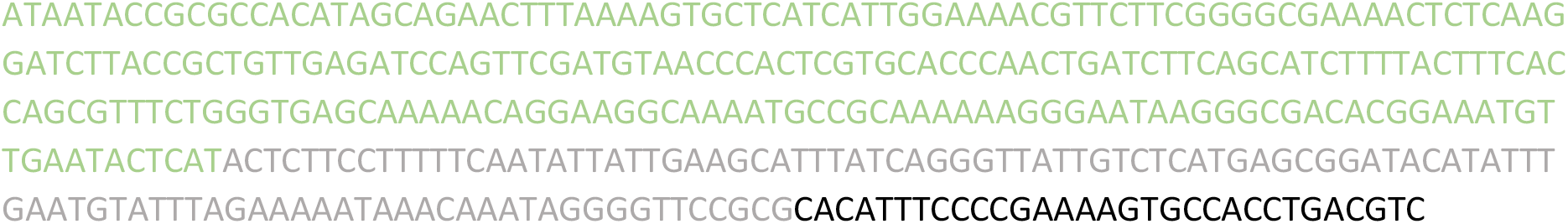
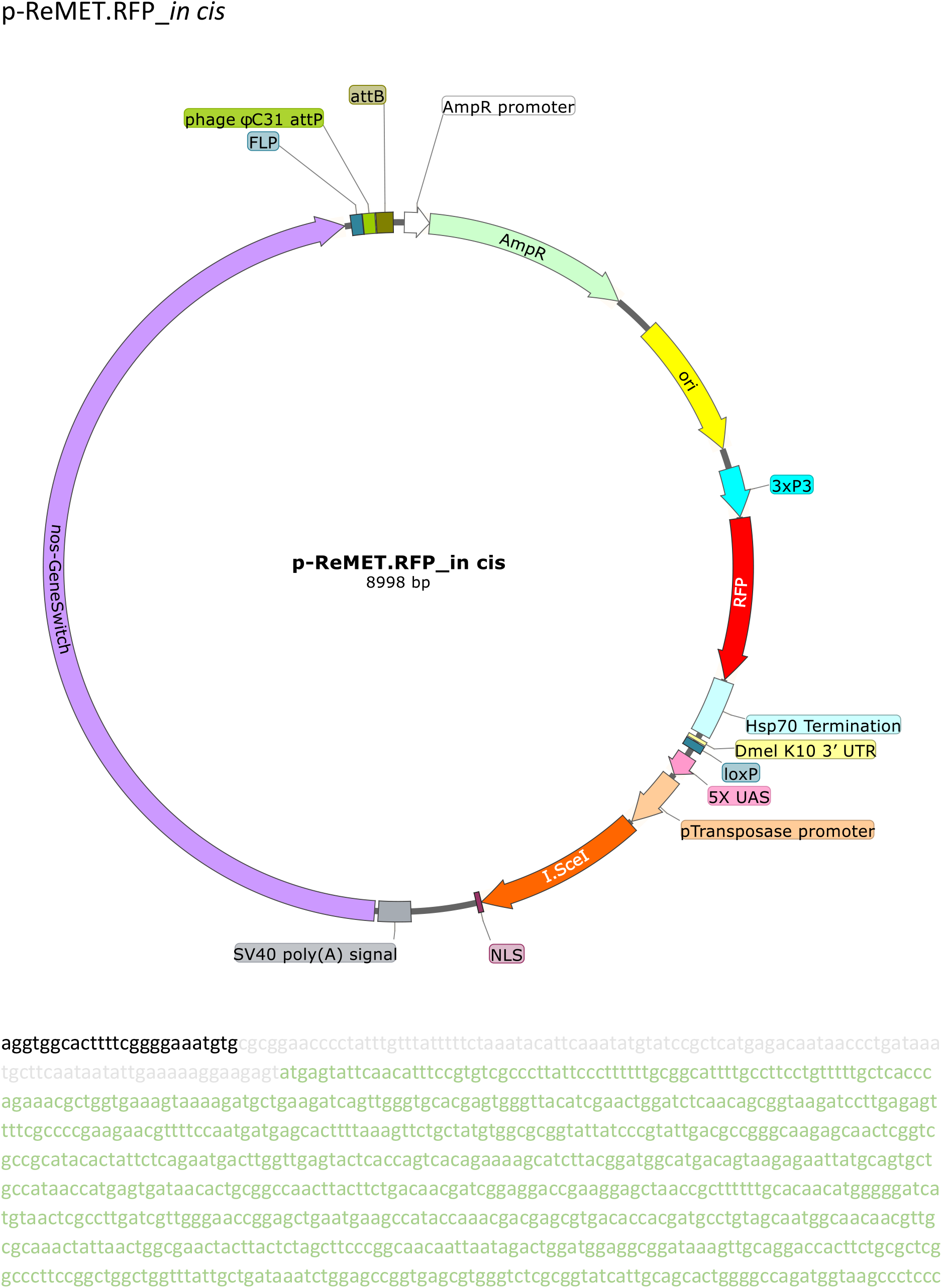
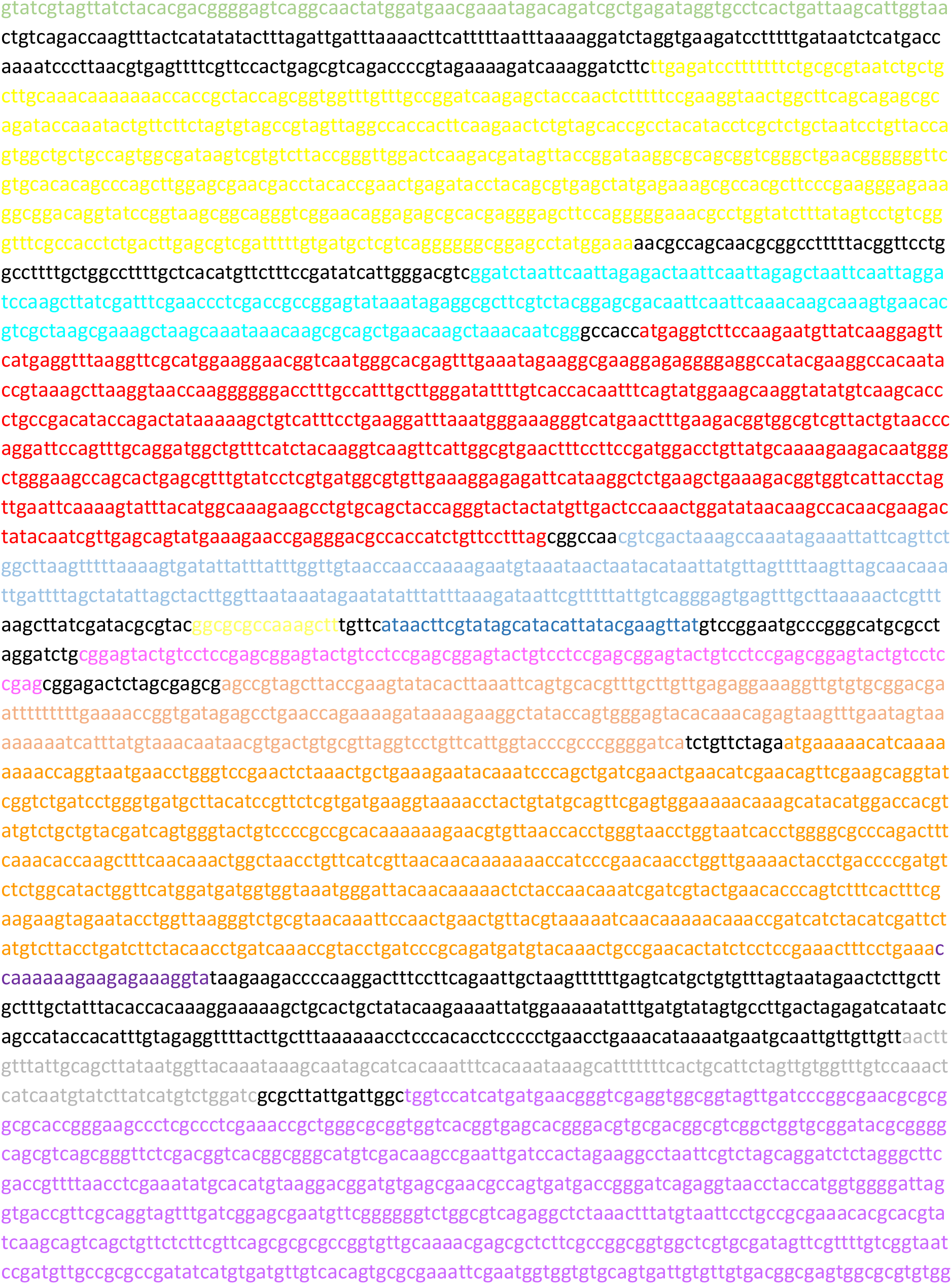
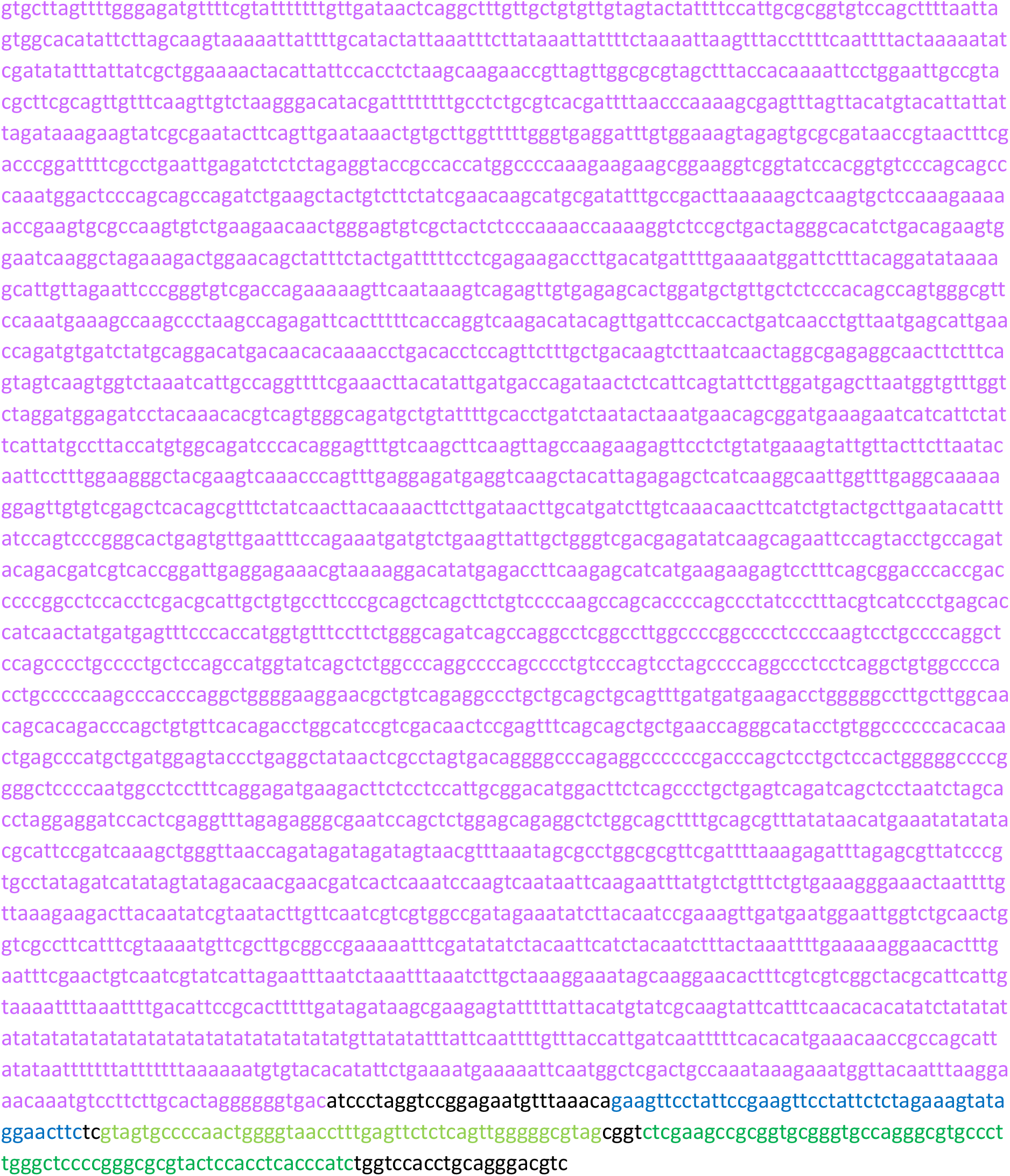
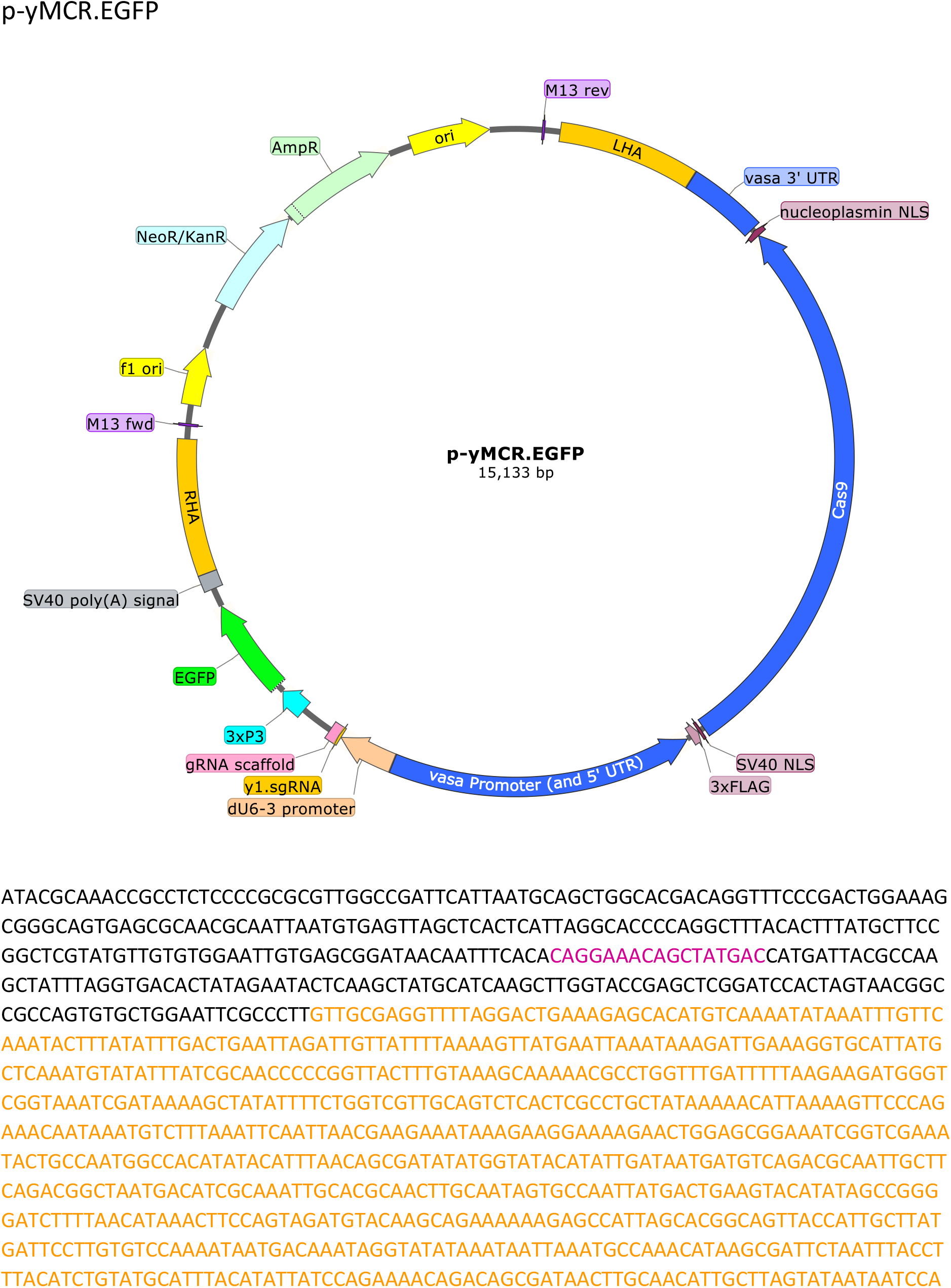
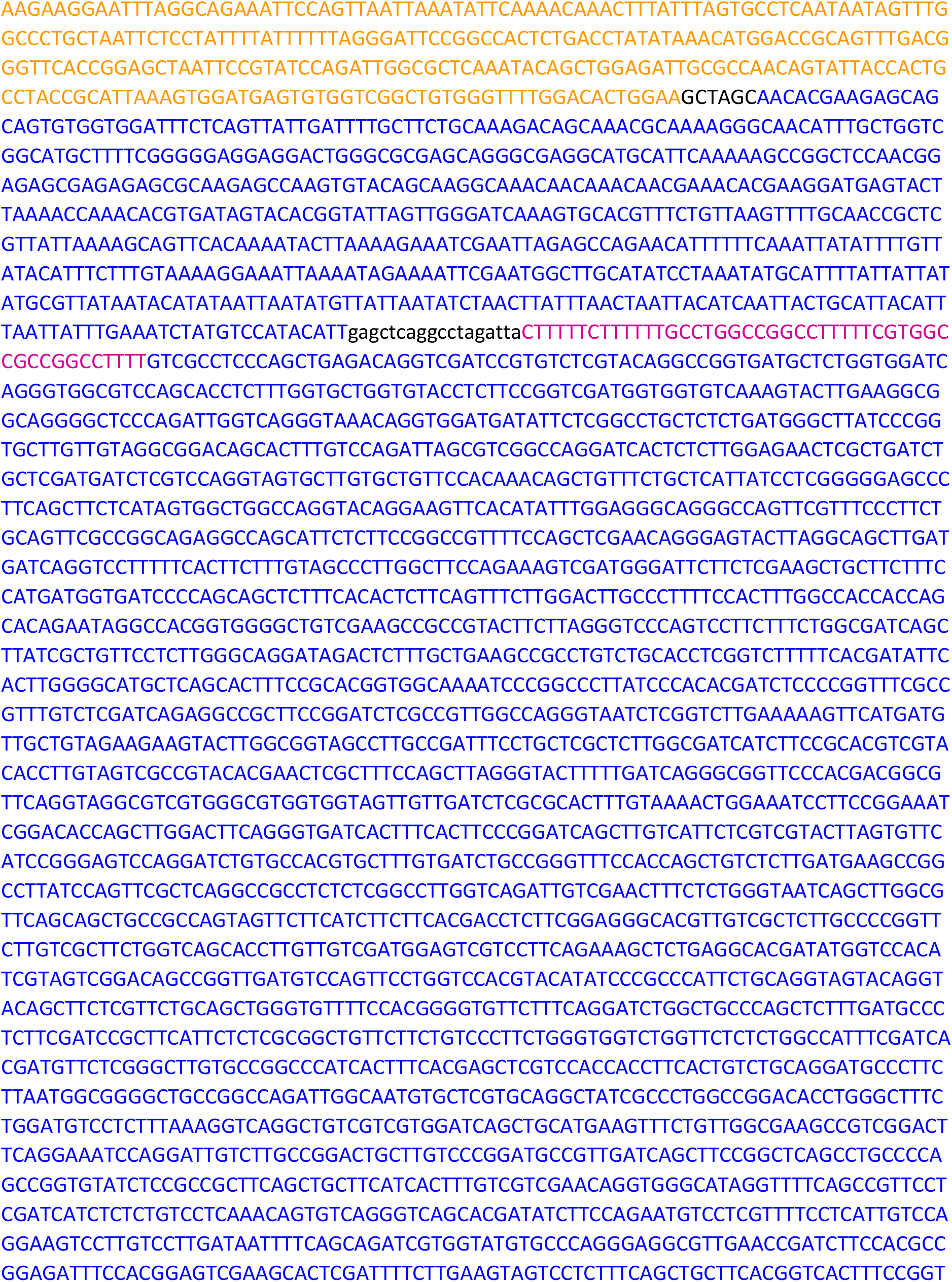
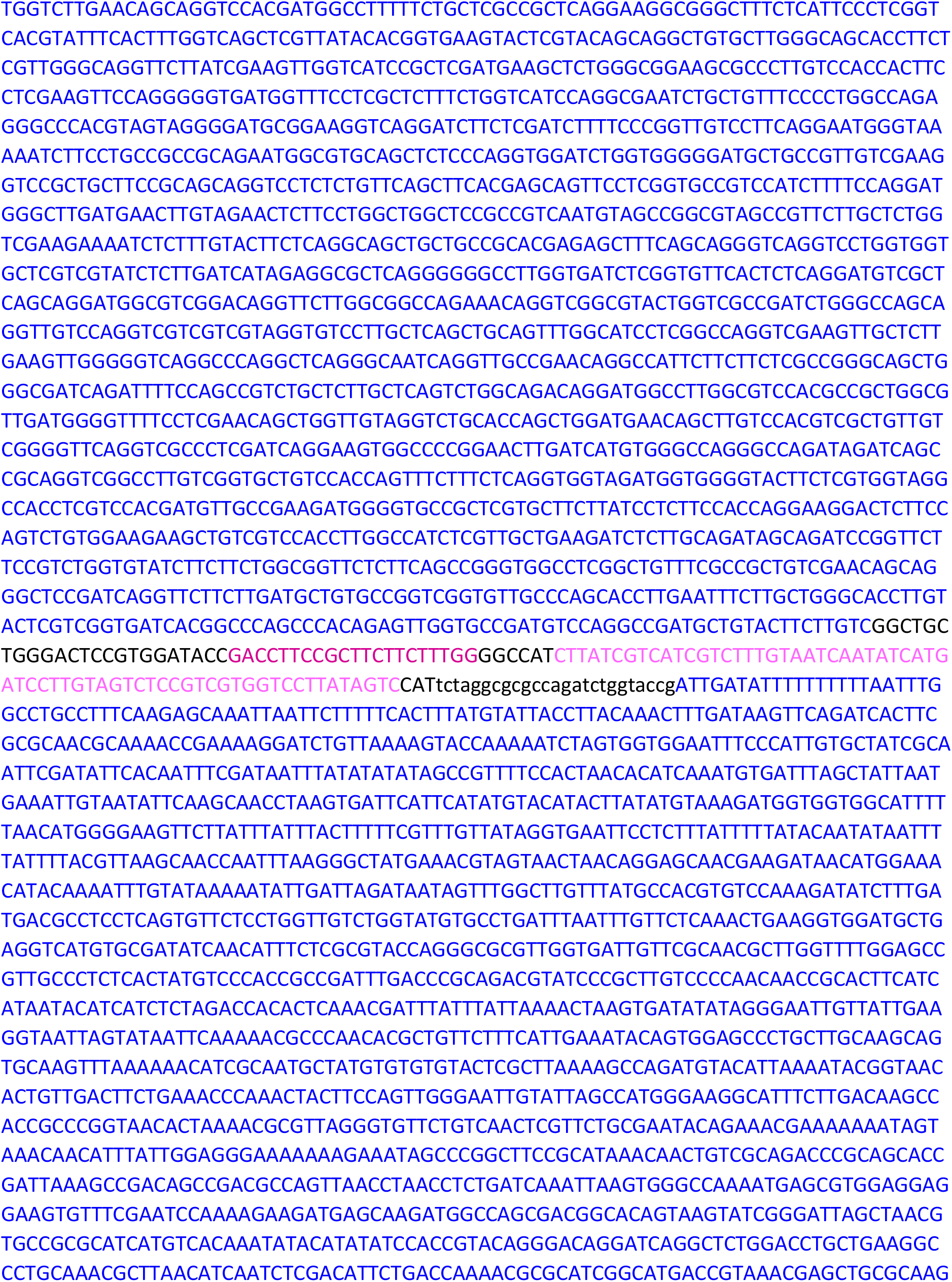
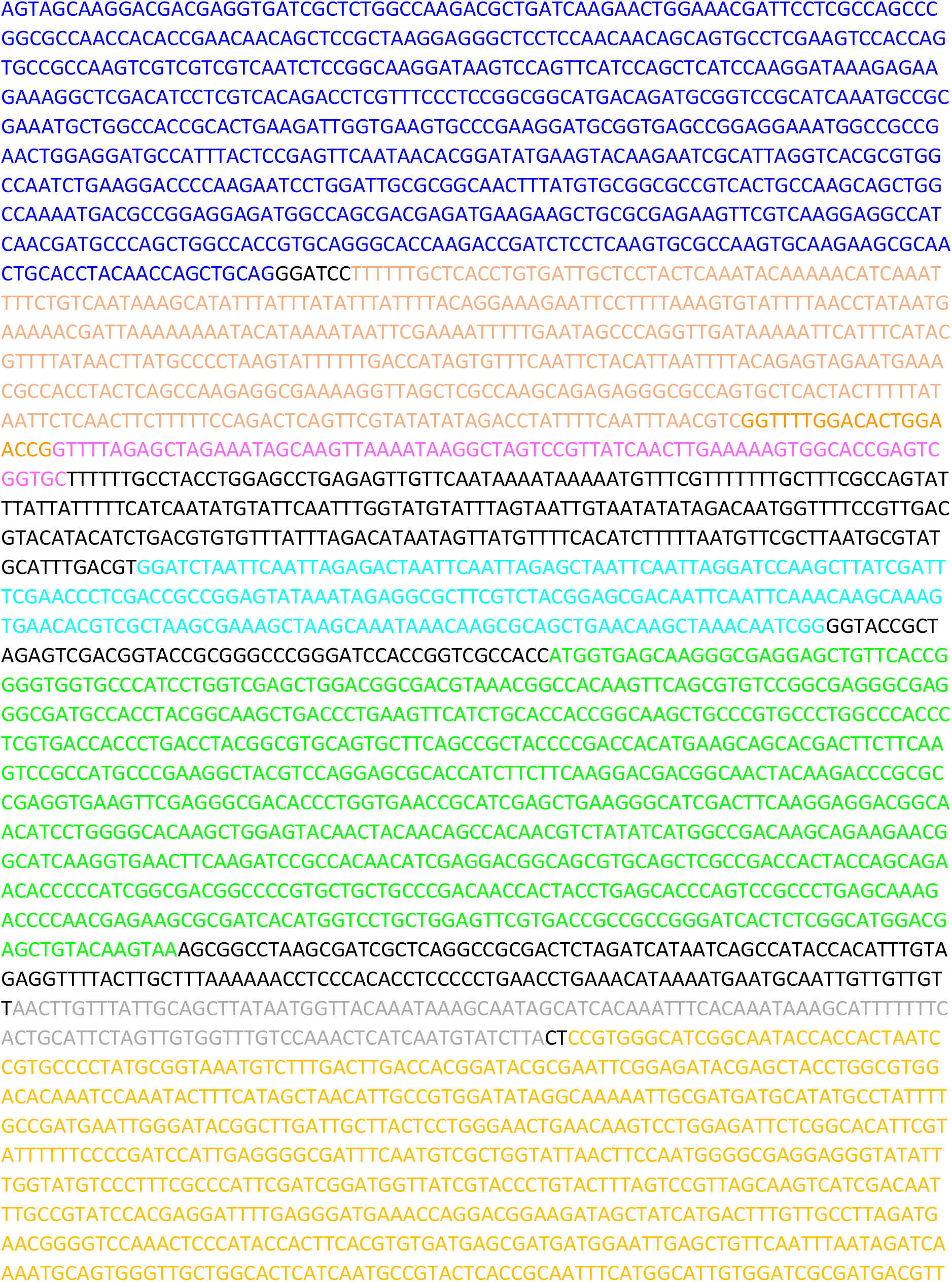
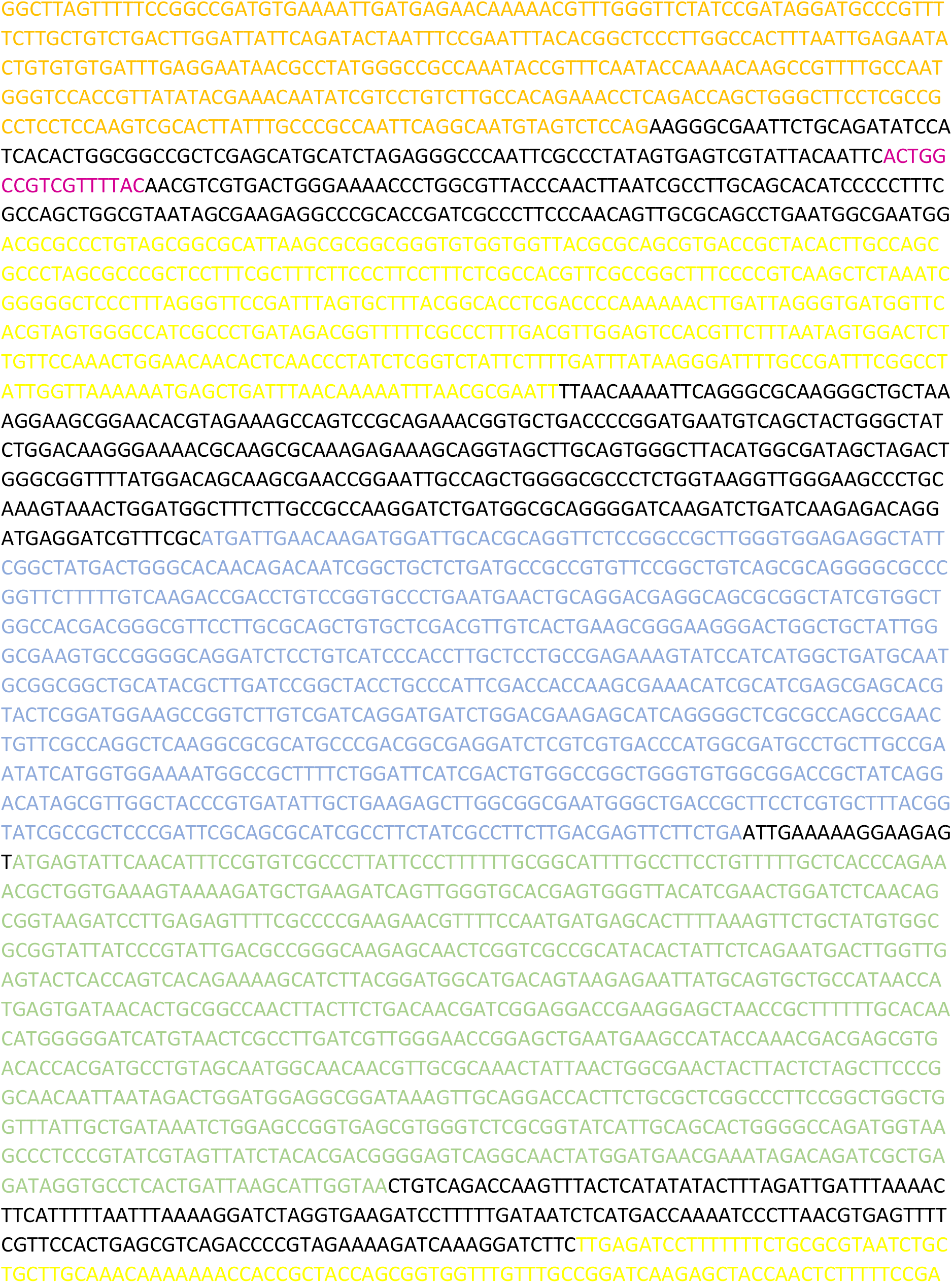
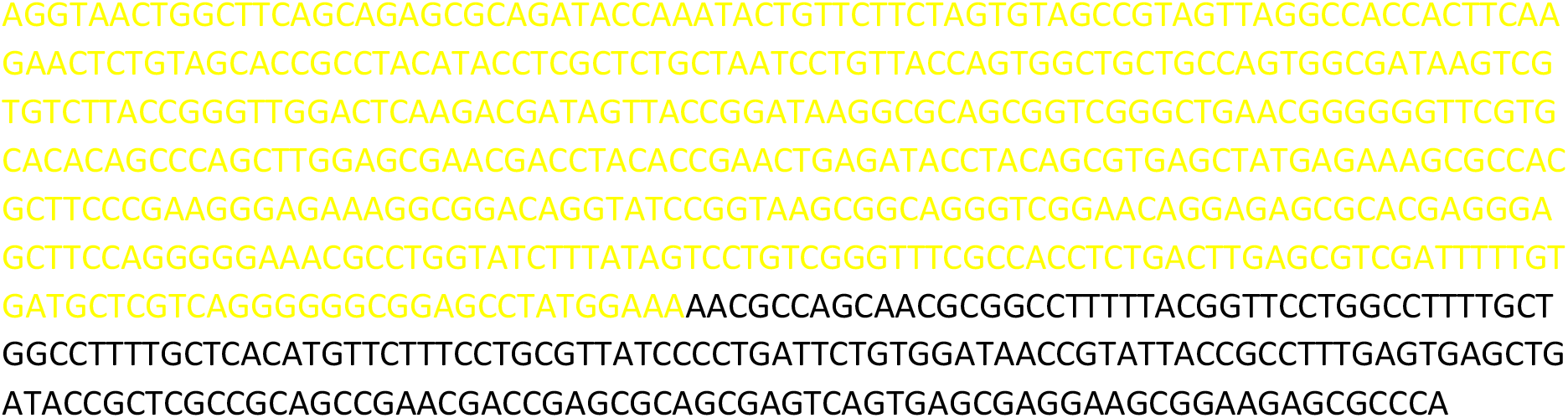
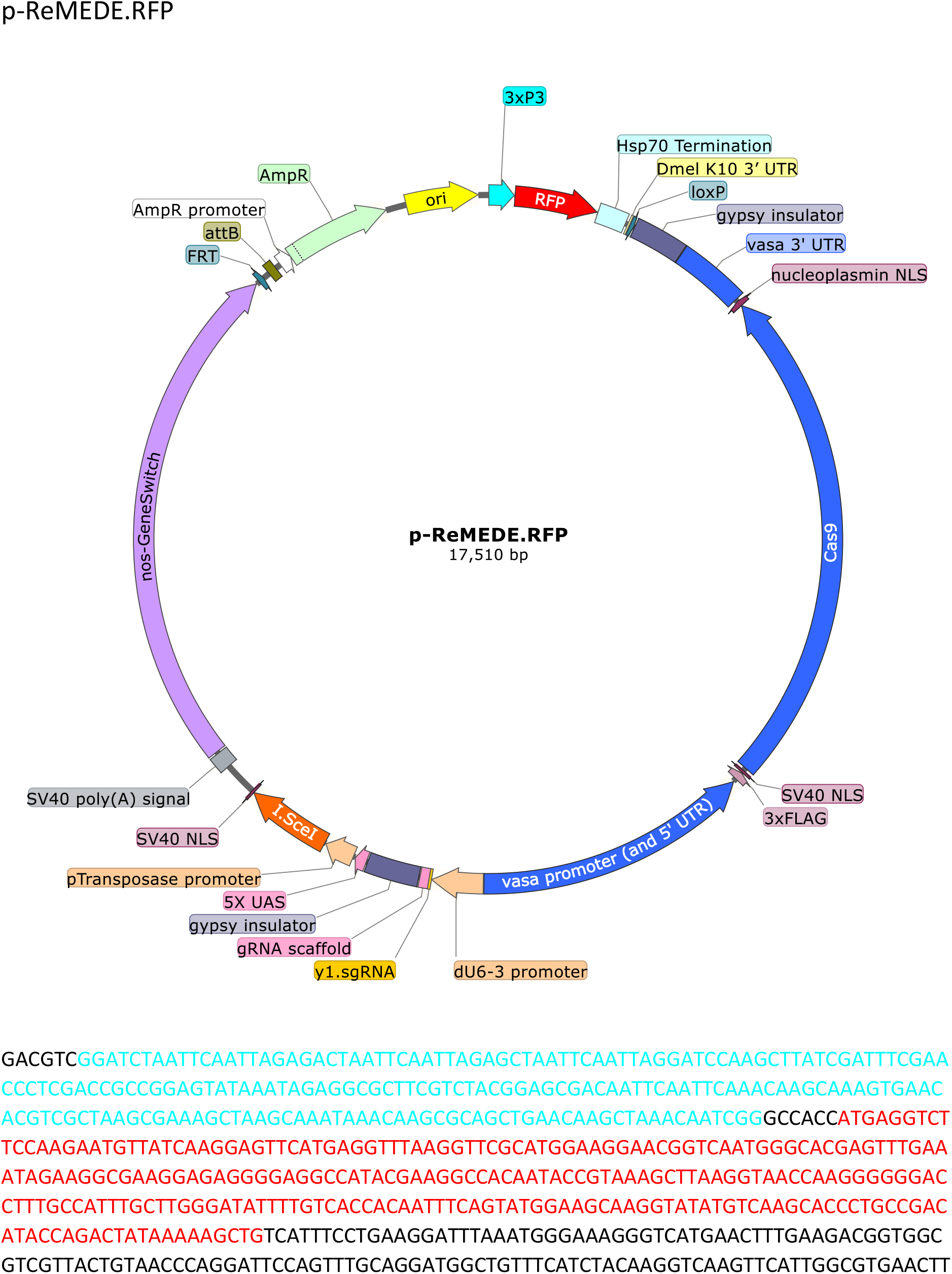
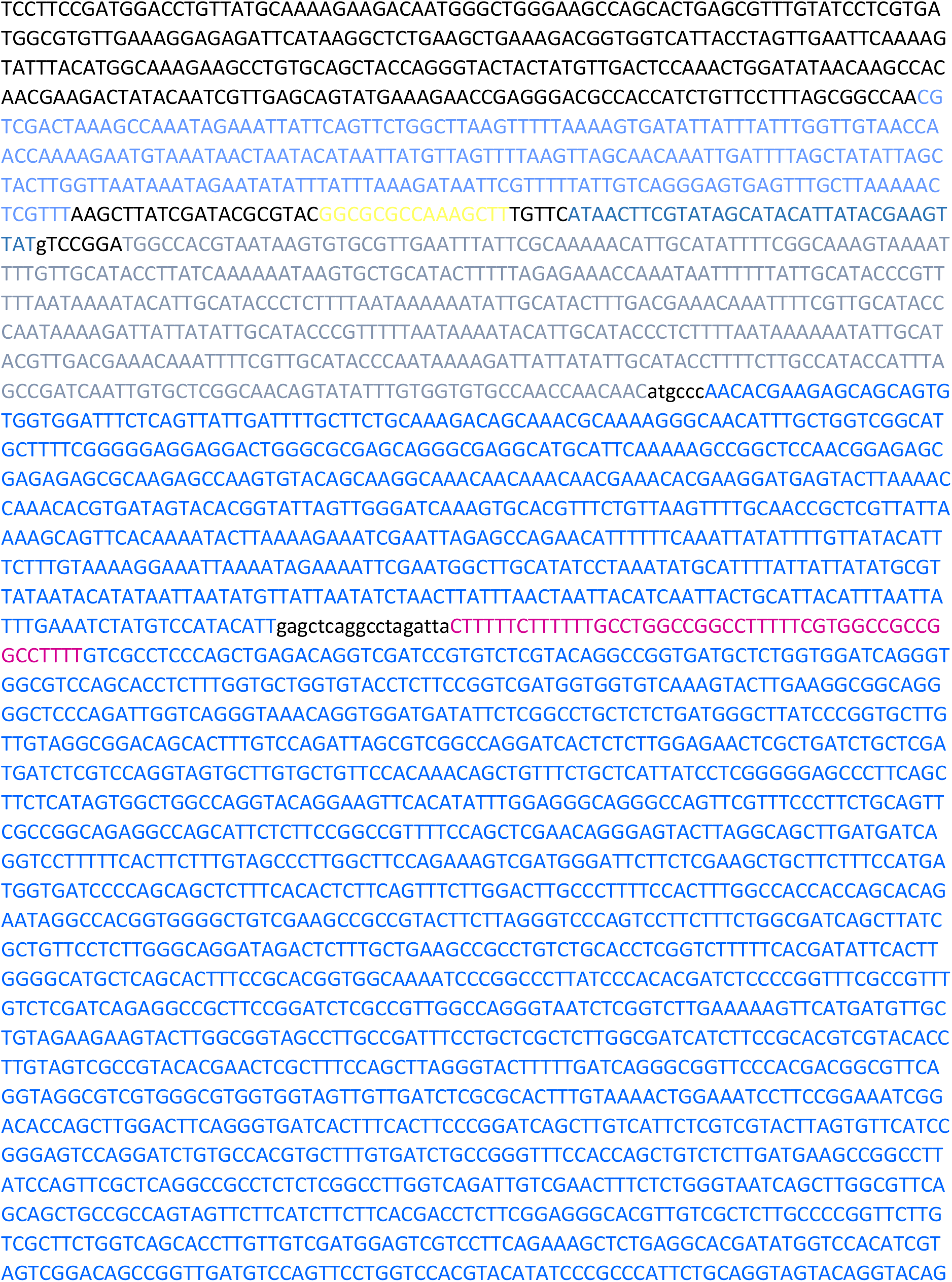
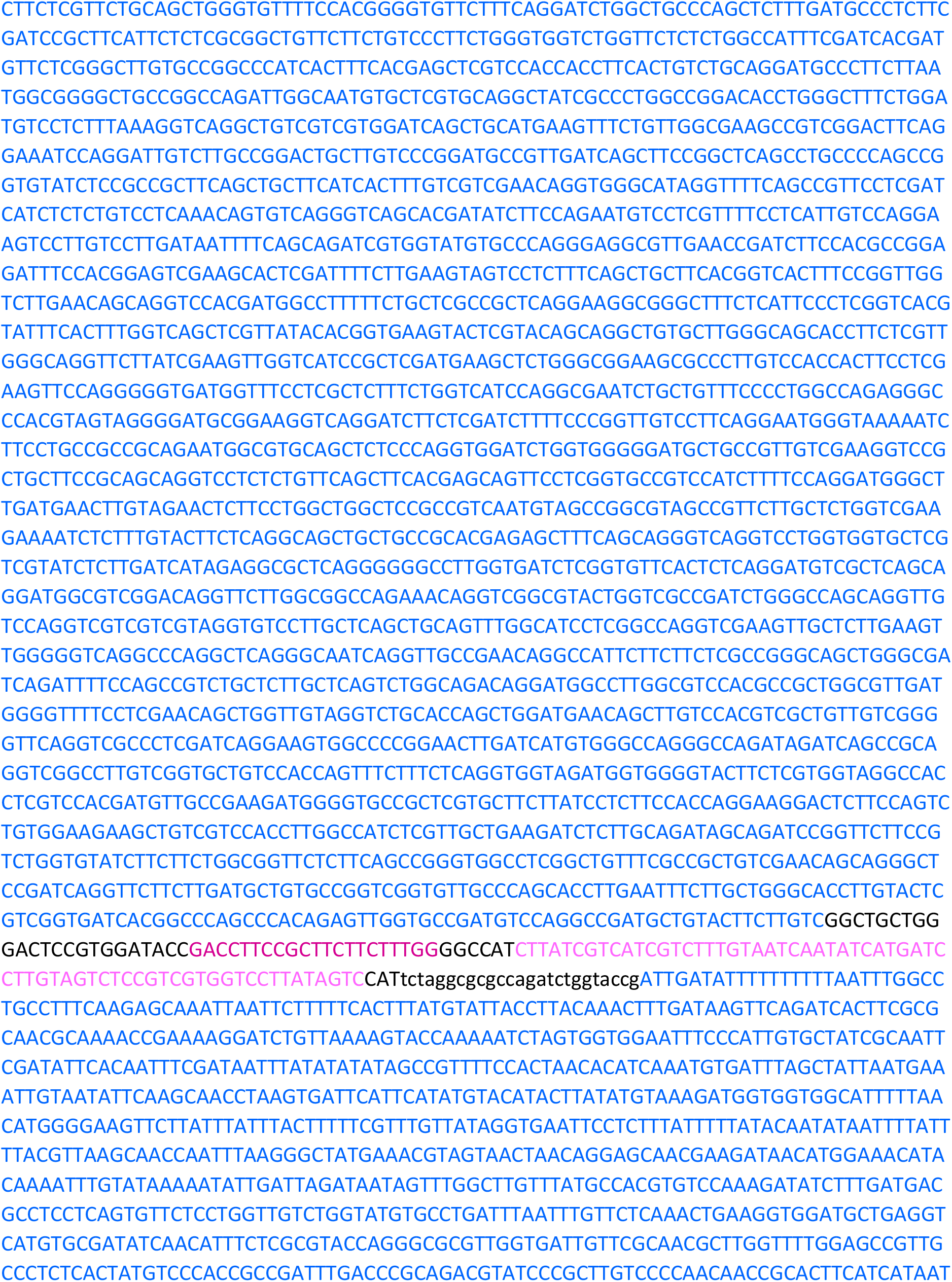
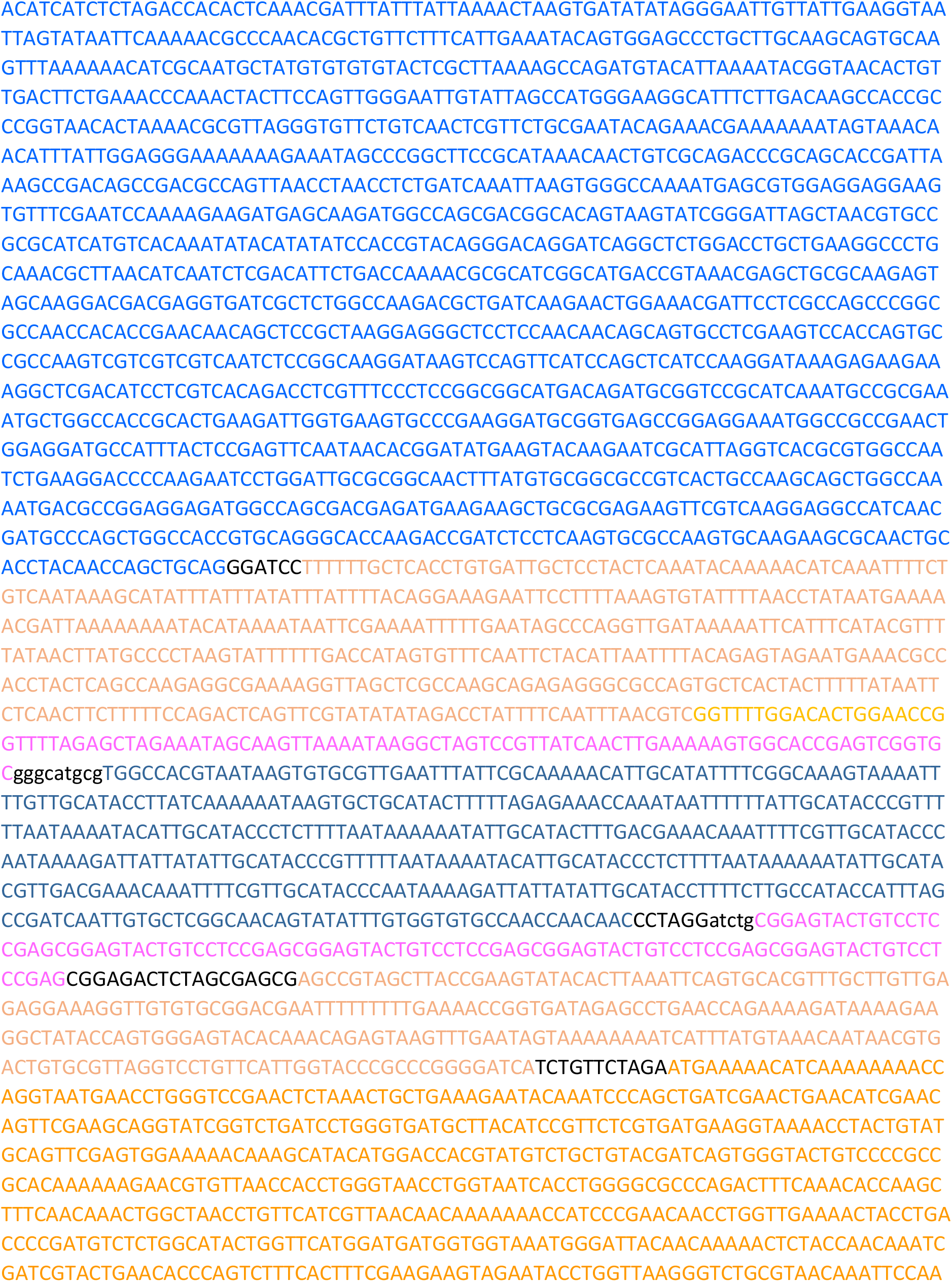
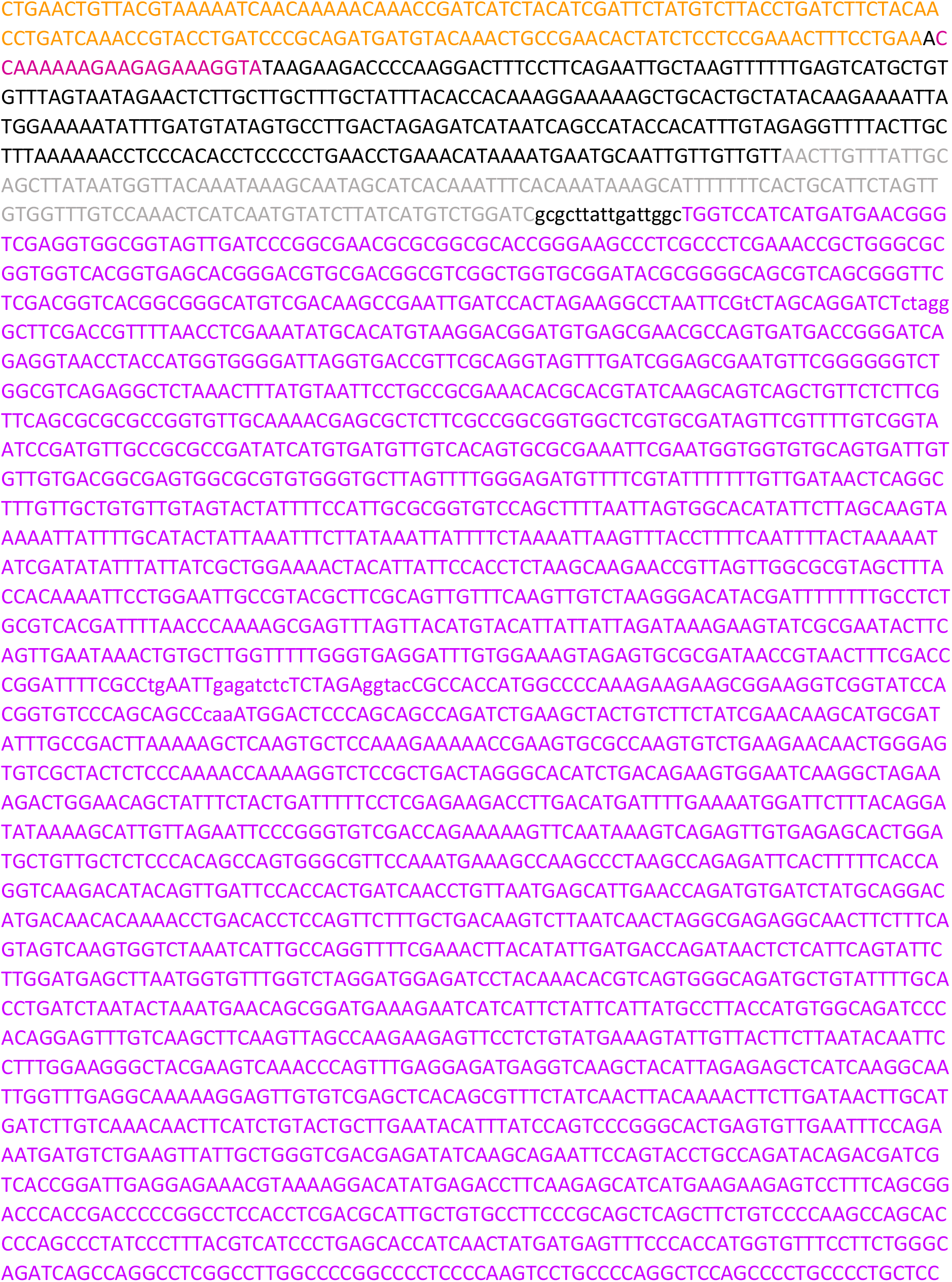
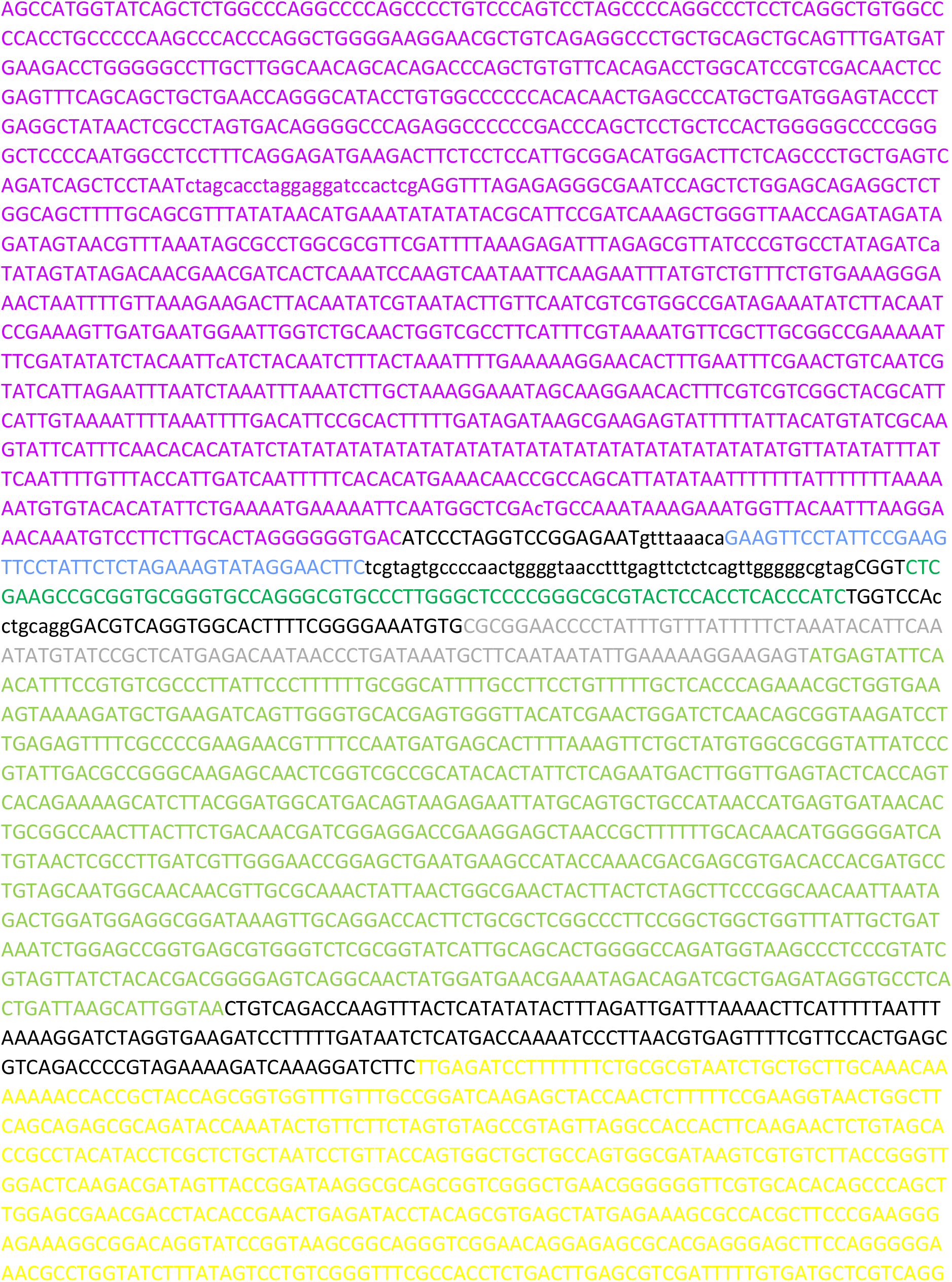
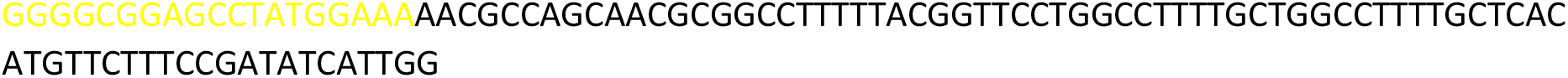
Plasmid Maps and Sequences.

